# EpiAgent: Foundation model for single-cell epigenomic data

**DOI:** 10.1101/2024.12.19.629312

**Authors:** Xiaoyang Chen, Keyi Li, Xuejian Cui, Zian Wang, Qun Jiang, Jiacheng Lin, Zhen Li, Zijing Gao, Rui Jiang

## Abstract

Large-scale foundation models have recently opened new avenues for artificial general intelligence. Such a research paradigm has recently shown considerable promise in the analysis of single-cell sequencing data, while to date, efforts have centered on transcriptome. In contrast to gene expression, chromatin accessibility provides more decisive insights into cell states, shaping the chromatin regulatory landscapes that control transcription in distinct cell types. Yet, challenges also persist due to the abundance of features, high data sparsity, and the quasi-binary nature of these data. Here, we introduce EpiAgent, the first foundation model for single-cell epigenomic data, pretrained on a large-scale Human-scATAC-Corpus comprising approximately 5 million cells and 35 billion tokens. EpiAgent encodes chromatin accessibility patterns of cells as concise “cell sentences,” and employs bidirectional attention to capture cellular heterogeneity behind regulatory networks. With comprehensive benchmarks, we demonstrate that EpiAgent excels in typical downstream tasks, including unsupervised feature extraction, supervised cell annotation, and data imputation. By incorporating external embeddings, EpiAgent facilitates the prediction of cellular responses to both out-of-sample stimulated and unseen genetic perturbations, as well as reference data integration and query data mapping. By simulating the knockout of key *cis*-regulatory elements, EpiAgent enables in-silico treatment for cancer analysis. We further extended zero-shot capabilities of EpiAgent, allowing direct cell type annotation on newly sequenced datasets without additional training.

## Introduction

Single-cell assay for transposase-accessible chromatin using sequencing (scATAC-seq) captures chromatin dynamics and reveals regulatory landscapes at the level of individual cells^1,2^, enabling the detection of cell heterogeneity^3^, tissue development^4^, and disease mechanisms^5^. With advances in sequencing technologies, a multitude of large-scale cell atlases that encompass fetal development^6^, adult tissues^7^, brain tissues^8^, and neural development^9^ have been constructed, providing unprecedented resources to uncover regulatory patterns under diverse physiological conditions. However, the vast number of candidate cis-regulatory elements (cCREs), the extreme sparsity, and its quasi-binary nature pose significant challenges for scATAC-seq data analysis.

Although numerous comprehensive pipelines^10–13^ and methods targeting specific tasks, such as unsupervised feature extraction^14–18^, supervised cell annotation^19–21^, and data imputation^22,23^, have been developed for scATAC-seq analysis, they still remain limited. First, most methods lack generalizability, being confined to solving single tasks with limited extensibility and requiring major modifications to address new tasks. Second, they fail to effectively leverage the latent cellular heterogeneity and regulatory network information embedded in large-scale atlas-level datasets, often requiring retraining from scratch for new datasets and leading to unstable performance across different datasets. Third, they use high-dimensional cell-by-cCRE matrices as input, necessitating prior filtering that excludes many accessible cCREs, thereby compromising model performance. Finally, key challenges remain unaddressed, such as predicting cellular responses to external stimulation or genetic perturbations and simulating cell state changes following cCRE knockouts. Consequently, there is a pressing need for the foundation model and innovative pretraining strategies tailored to the unique characteristics of scATAC-seq data.

Pretrained foundation models have emerged as a powerful paradigm in natural language processing^24^, computer vision^25^, and multimodal learning^26^, and are now becoming integral to the pursuit of artificial general intelligence^27^. Within single-cell sequencing, foundation models^28–32^ have primarily been employed to simultaneously resolve cell representations and capture gene co-expression patterns in transcriptomic data^33,34^. In contrast, chromatin accessibility data contain more fundamental regulatory context that governs transcription, while developing a transformer-based foundation model to harness these data faces two major obstacles. First, the immense number of cCREs in total but few accessible in each cell, coupled with the values of quasi-binary nature, makes it challenging to formulate a cell sentence that meaningfully integrates these elements. Second, unlike words in a linguistic context, cCREs lack explicitly directional regulatory relationship, making traditional context-based pretraining strategies (e.g., masked language modeling or generative pretraining tasks) less directly applicable.

Here we proposed EpiAgent, a transformer-based foundation model for single-cell epigenomic data. For providing comprehensive pretraining resources for EpiAgent, we constructed a large-scale corpus, Human-scATAC-Corpus, comprising approximately 5 million cells and 35 billion accessible cCREs. For each cell, EpiAgent considers only its accessible cCREs, ordering them by importance to form a cell sentence as input. EpiAgent has approximately 1.4 billion parameters, including a cCRE embedding module, the EpiAgent transformer, and a signal decoder. EpiAgent is pretrained using a newly designed cell-cCRE alignment task, along with a signal reconstruction task, to learn fundamental patterns of cellular heterogeneity and regulatory networks by its bidirectional self-attention mechanism.

To comprehensively evaluate EpiAgent, we first fine-tuned it on three typical downstream tasks, including unsupervised feature extraction, supervised cell annotation, and data imputation, and demonstrated its superior performance. Notably, when the dataset contains cell populations similar to Human-scATAC-Corpus, EpiAgent can directly produce cell embeddings rich in heterogeneity in zero-shot conditions. By incorporating perturbation-specific token embedding into the head token embedding, EpiAgent can accurately predict cellular responses to both out-of-sample stimulated and unseen genetic perturbations. By substituting external information with batch-specific token embeddings, EpiAgent effectively reduces or even eliminates batch effects when integrating data as a cell atlas. Using the cell atlas as reference, EpiAgent enables query data mapping with followed label transfer, outperforming the analysis of the query data alone. Benefiting from the unique modeling of cell sentences, EpiAgent enables flexible in-silico cCRE knockouts in cancer cells, facilitating the quantitative evaluation of their effects on cell states. To further extend zero-shot capabilities to cell type annotation, we developed EpiAgent-B and EpiAgent-NT, for directly annotating cells from brain and other tissues, respectively, without requiring additional training. Together, EpiAgent represents a significant leap forward in single-cell epigenomic analysis, enabling more precise and comprehensive insights into cellular mechanisms.

## Results

### Overview of EpiAgent

Due to the inherent high sparsity and dimensionality of scATAC-seq data, EpiAgent focuses on tokenizing only the cCREs that are accessible in cells. The ranks of tokens in a cell tokenized sentence are determined by the values after term frequency-inverse document frequency (TF-IDF) transformation (Fig. 1b).

**Fig. 1.**
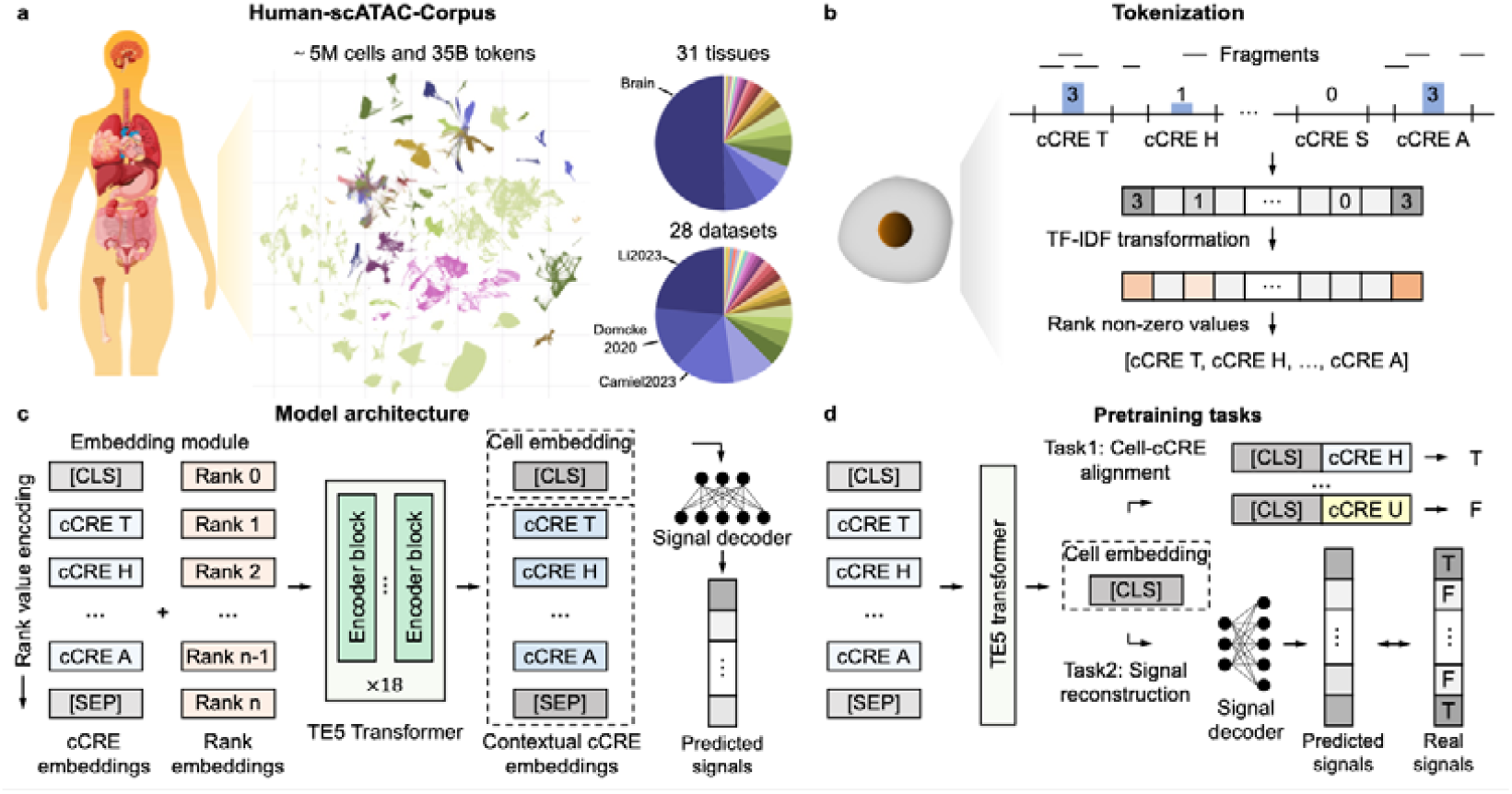
Overview of EpiAgent. **a,** Human-scATAC-Corpus, comprising approximately 5 million cells and 35 billion tokens from 31 tissues and 28 datasets, providing a valuable resource for pretraining EpiAgent. **b**, Tokenization in EpiAgent, where accessible cCREs in scATAC-seq data are ranked in descending order based on their TF-IDF-transformed values to form cell sentences. **c**, Model architecture of EpiAgent, consisting of three modules: (1) an embedding module, that converts cell sentences into embedding sequences, (2) the EpiAgent transformer, with multiple transformer blocks using bidirectional attention mechanisms to generate cell embeddings and contextual cCRE embeddings, and (3) a signal decoder that reconstructs signals from cell embeddings. **d**, Pretraining tasks for EpiAgent, including the cell-cCRE alignment task and the signal reconstruction task. EpiAgent is a transformer-based foundation model pretrained on our custom-built, large-scale scATAC-seq corpus, Human-scATAC-Corpus (Fig. 1a). Human-scATAC-Corpus comprises data from 31 different tissues, 28 public datasets, approximately 5 million cells, and 35 billion tokens, providing EpiAgent with diverse and comprehensive resources to learn general epigenetic regulatory patterns across various tissues and cell types during unsupervised pretraining.

EpiAgent consists of three modules: an embedding module (∼695 million parameters), the EpiAgent transformer (∼57 million parameters), and a signal decoder (∼695 million parameters), bringing the total parameter count to approximately 1.4 billion (Fig. 1c). A tokenized cell sentence is augmented with two special tokens, [CLS] (the classification token) and [SEP] (the separator token), at the beginning and end, respectively. The embedding module then converts the sentence into cCRE embeddings and rank embeddings, analogous to word embeddings and positional embeddings in natural language processing. The EpiAgent transformer takes the sum of the two sets of embeddings as input, and uses the bidirectional attention mechanism to capture cCRE co-accessibility patterns, which can be regarded as regulatory networks at the chromatin level. The EpiAgent transformer consists of 18 layers of transformer encoder blocks, and outputs contextualized cCRE embeddings that corresponds directly to the input cCRE embeddings. Flash Attention v2^35^ is also used for accelerating steps for training and inference. The output [CLS] embedding represents the cell embedding, and additional information can be introduced by adding a conditional embedding to the input [CLS] token embedding, allowing for in-silico simulation of changes in cell states. Finally, the signal decoder, consisting of a single-layer fully connected neural network, reconstructs the accessibility signal of all cCREs for each cell from the cell embeddings.

The pretraining tasks for EpiAgent are designed to simultaneously capture cCRE co-accessibility patterns and generate comprehensive cell embeddings, encompassing two key tasks: the cell-cCRE alignment task and the signal reconstruction task (Fig. 1d). In the cell-cCRE alignment task, EpiAgent concatenates the output cell embedding with the input embeddings of cCREs that are either accessible or inaccessible in the cell, and performs a binary classification to determine their accessibility. In the signal reconstruction task, EpiAgent directly feeds the output cell embedding into the signal decoder to reconstruct the accessibility signals for all cCREs in the cell. After undergoing full pretraining on the Human-scATAC-Corpus, EpiAgent effectively captures cellular heterogeneity and regulatory networks across diverse tissues, sequencing technologies, and sequencing depths, achieving superior performance across a range of downstream tasks. The details regarding the Human-scATAC-Corpus, tokenization strategy, model architecture, and pretraining tasks are available in Methods.

### EpiAgent excels in typical downstream tasks of scATAC-seq data analysis

Typical downstream analyses of single-cell data, including unsupervised feature extraction, supervised cell annotation, and data imputation, rely on accurate and robust computational frameworks^36^. As a foundation model for scATAC-seq data, EpiAgent serves as a core module that can be fine-tuned and integrated with task-specific modules to adapt to various downstream tasks (Fig. 2a). To comprehensively benchmark EpiAgent, we collected four additional datasets varying in similarity to Human-scATAC-Corpus: Buenrostro2018^37^ (hematopoietic cell differentiation), Kanemaru2023^38^ (heart), Li2023b^39^ (brain), and Ameen2022^40^ (stem cells cultured in vitro) datasets. Among these, Kanemaru2023 and Li2023b datasets, derived from human tissues, have similar cell populations with data in the Human-scATAC-Corpus, while Buenrostro2018 and Ameen2022 datasets contain cell types that differ significantly from those in the pretraining corpus.

**Fig. 2.**
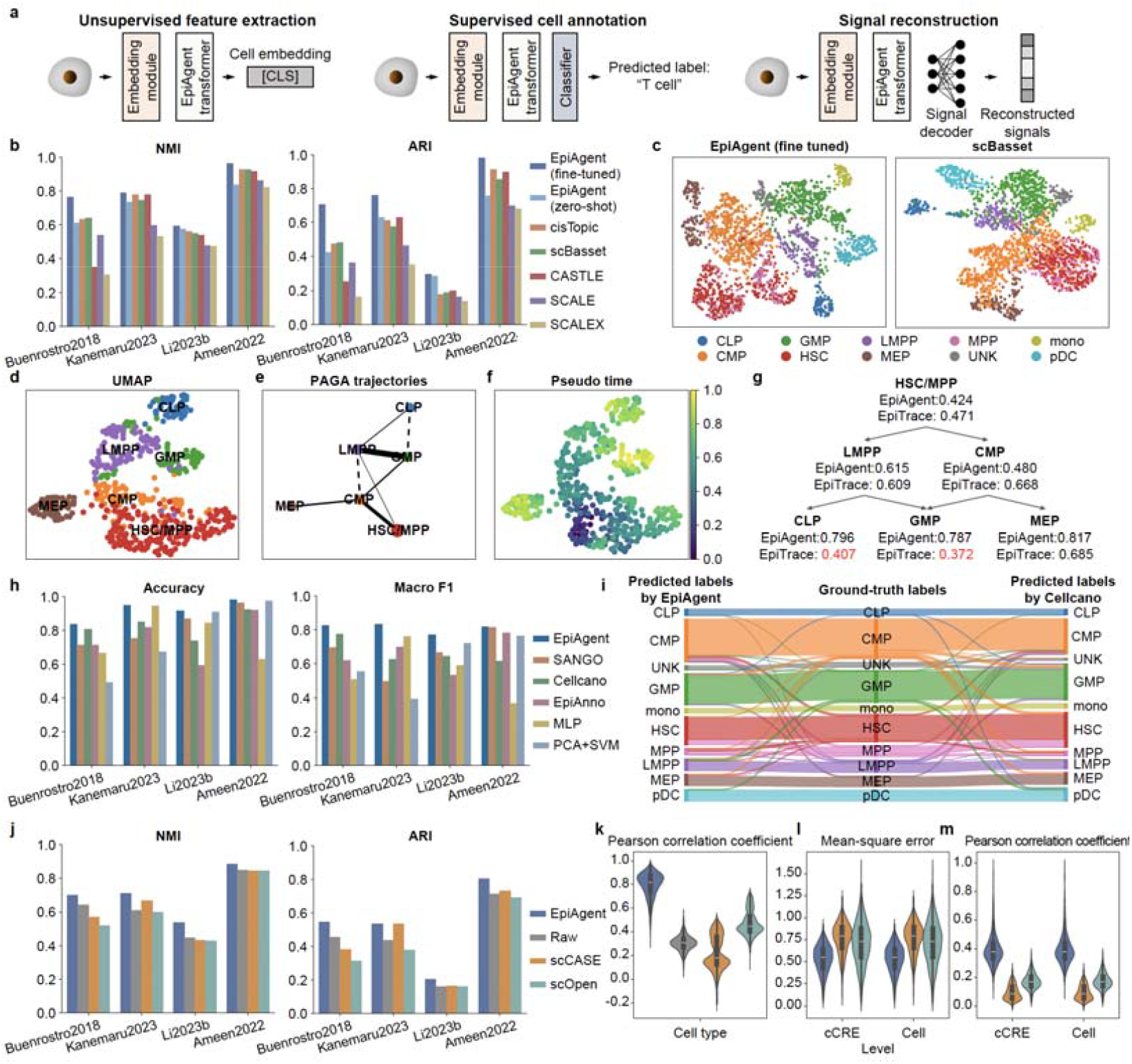
Typical downstream tasks of scATAC-seq data with EpiAgent. **a,** Overview of EpiAgent fine-tuned for various downstream tasks, including unsupervised feature extraction, supervised cell annotation, and data imputation. **b**, Benchmarking performance on unsupervised feature extraction evaluated by NMI and ARI scores. **c**, UMAP visualization of cell embeddings derived from the fine-tuned EpiAgent model compared to the second-best model, scBasset, on the Buenrostro2018 dataset. **d-g**, Analysis of donor “BM0828” cells in the Buenrostro2018 dataset, including UMAP visualization of cell embeddings (**d**), PAGA trajectories (**e**), pseudo-time estimation (**f**), and the average pseudo-time of each cell type (**g**). UMAP visualizations (**d-f**), PAGA trajectories (**e**), and pseudo-time estimations (**e & f**) are derived from cell embeddings obtained using the fine-tuned EpiAgent model. Dashed lines in **e** indicate trajectories between sibling cell types. Pseudo-time (**e & f**) incorporates additional ClockDML information. **h**, Benchmarking performance on supervised cell type annotation evaluated by accuracy and macro F1 scores. **i**, Sankey plot among EpiAgent predicted labels, ground-truth labels and Cellcano (the second-best method) predicted labels on the Buenrostro2018 dataset. **j**, Benchmarking performance of data imputation evaluated by clustering metrics (NMI and ARI scores), and clustering is performed using PCA on imputed matrices from each method. **k**, Pearson correlation coefficients between imputed signals from each method and average raw signals of the corresponding cell types. **l & m**, Mean-square errors (**l**) and Pearson correlation coefficients (**m**) between imputed signals from each method and raw signals of their corresponding cell types for each method. In **k-m**, the upper, middle, and lower edges of the boxes denote the upper quartile, median, and lower quartile, respectively, and error bars denote the maximum and minimum values.

#### Unsupervised feature extraction

Given the high dimensionality and extreme sparsity of scATAC-seq data, direct analysis in the raw data space is infeasible, thus reducing scATAC-seq data to low-dimensional cell embeddings through feature extraction is a critical step. Using a strategy consistent with pretraining, we fine-tuned EpiAgent on new datasets and compared its zero-shot and fine-tuned performance against five baseline methods: cisTopic^14^, scBasset^17^, CASTLE^18^, SCALE^15^, and SCALEX^16^, with implementations followed open-source codes and default configurations. Notably, scBasset additionally incorporates cCRE sequence information, while other baselines rely solely on peak-by-cell matrices. Cell embeddings are clustered using the Leiden algorithm, with the number of clusters aligned to the known cell types via a binary search approach. Clustering performance is evaluated using normalized mutual information (NMI) and adjusted Rand Index (ARI) scores, where higher values indicate embeddings that better capture cellular heterogeneity. Details on experimental setup, metric calculations, fine-tuning of EpiAgent, and baseline methods are provided in Methods.

As shown in Fig. 2b, the fine-tuned EpiAgent outperforms all baselines, with an average improvement of 7.214% and 25.975% in NMI and ARI, respectively, compared to the second-best model, cisTopic. On datasets with cell populations similar to those in the Human-scATAC-Corpus, EpiAgent performs competitively with baseline methods even without fine-tuning, particularly on the Li2023b dataset, owing to the brain cells constituting a substantial proportion of the pretraining data. UMAP visualizations (Fig. 2c and Supplementary Fig. 1-4) further illustrate that EpiAgent better separates distinct cell types and compresses intra-type variability, highlighting its ability to capture cellular heterogeneity. Such superiority stems from not only unique attention mechanism and pretraining strategy in the EpiAgent transformer to extract intrinsic information of cells, but also its modeling of cell sentences to focus on the widest possible range of accessible cCREs. In contrast, baseline methods relying on cell-by-cCRE matrices are inherently limited by extremely sparse signals and filtered cCREs.

The cell embeddings derived from EpiAgent also reveal clear developmental trajectories. We first used the fine-tuned EpiAgent model to generate embeddings for cells of donor “BM0828” in the Buenrostro2018 dataset (Fig. 2g), and then performed PAGA to infer trajectories (Fig. 2e). Except for a few trajectories between sibling cell types (indicated by dashed lines), the inferred trajectories align well with hematopoietic differentiation. Furthermore, we combined clock-like differential methylation loci (ClockDML) identified by Xiao et al.^41^ with these embeddings to infer cellular pseudotime, which also can be served as relative mitotic age, without relying on inferred trajectories (Supplementary Note 1). As shown in Fig. 2f and Fig. 2g, EpiAgent accurately predicts pseudotime for hematopoietic differentiation, aligning well with the differentiation trajectories, while in contrast, the EpiTrace model^41^ mispredicts the ages of CLP and GMP cells. All the results above demonstrate that EpiAgent not only delineates discrete cell types but also captures developmental heterogeneity effectively.

#### Supervised cell annotation

Supervised cell annotation leverages reference dataset labels to train models for identifying cell types in query data, which is another crucial task in scATAC-seq analysis. For fine-tuning, we fed the EpiAgent output [CLS] token embeddings into a multilayer perceptron (MLP) classifier to adapt for this task. We compared EpiAgent against five baseline methods: Principal Component Analysis and Support Vector Machine (PCA+SVM)^42^, MLP^43^, EpiAnno^19^, CellCano^20^, and SANGO^21^. Although PCA+SVM and MLP are not specifically designed for scATAC-seq data, they are selected due to their superior performance in benchmarks for methods based on scRNA-seq data^42,43^. Each dataset is randomly split, with two-thirds serving as the reference dataset for training and the remaining one-third as the query dataset. Annotation performance is evaluated by accuracy and macro F1, with macro F1 revealing the ability of models for annotating rare cell types. Details on experimental setup, metric calculations, fine-tuning of EpiAgent, and baseline methods are provided in Methods.

As shown in Fig. 2h, EpiAgent achieves the superior performance across datasets, outperforming the second-best methods by an average of 11.036% in accuracy (CellCano) and 21.549% in macro F1 (SANGO). Sankey diagrams (Fig. 2i) and confusion matrices (Supplementary Figs. 4-8) also highlight the robust annotation of EpiAgent across diverse cell population distributions or dataset sizes, accurately identifying both major and rare cell types. Such the stability can be attributed to the unique pretraining strategy for EpiAgent, while the performance of baseline methods shows considerable variability across datasets, suggesting their limited generalizability.

#### Data imputation

The extreme sparsity and low signal-to-noise ratio of scATAC-seq data can introduce substantial bias in downstream analyses, therefore data imputation is also a crucial step. Leveraging its signal decoder, EpiAgent can reconstruct cell-by-cCRE matrices from cell embeddings. We compared EpiAgent with raw data and two imputation methods specifically designed for scATAC-seq, scCASE^23^ and scOpen^22^, following the same evaluation that contains a unified dimensionality reduction and clustering as in scCASE and scOpen. Notably, scCASE and scOpen use complete cell-by-cCRE matrices as input, whereas EpiAgent only requires cell sentences composed of accessible cCRE indices. Detailed experimental settings, metric calculations, EpiAgent fine-tuning, and baseline descriptions are provided in Methods.

Clustering metrics reveal that EpiAgent achieved stable and efficient data imputation, with imputed data showing an average improvement of 11.123% in NMI and 18.605% in ARI over raw data. In contrast, baseline methods get inconsistent performance, sometimes underperforming compared to direct analysis of raw signals. UMAP visualizations also support the clustering results (Supplementary Fig. 9-12). To further validate imputation quality, we averaged signals from cells of the same cell type as ground truths and compared them to imputed signals using Pearson correlation coefficients. EpiAgent achieves the highest correlation, with median values exceeding 0.8 (Fig. 2m). Although EpiAgent does not explicitly aggregate embeddings based on cell similarity in latent space, as scCASE and scOpen do, its transformer architecture effectively enhances cellular heterogeneity and filters out irrelevant noise when reconstructing signals. Moreover, compared to baseline methods, EpiAgent consistently achieves lower MSE values and higher Pearson correlation coefficients between imputed and raw data at both the cCRE and cell levels, suggesting the ability to effectively reconstruct raw signals from compressed cell sentences.

### EpiAgent accurately predicts cellular responses to out-of-sample stimulated and unseen genetic perturbations

Single-cell characterization of cellular responses to perturbations illuminates fundamental principles of gene regulation and cellular function^44^, thereby informing pharmacological strategies, clinical interventions, and diagnostic advances^45,46^. Perturbations generally fall into two categories: external stimulations, such as electric shocks or drug treatments, and genetic perturbations, such as CRISPR-mediated gene editing. Advances in single-cell sequencing technologies have enabled the profiling cellular responses to these perturbations at the epigenomic level. However, due to the challenges posed by the high dimensionality and sparsity of scATAC-seq data, no existing methods have been specifically developed for single-cell perturbation prediction at the epigenomic level. Leveraging the transformer-based architecture, EpiAgent introduces an innovative approach by modifying cell sentences: it incorporates a perturbation-specific token into the [CLS] token of unperturbed cells to predict their post-perturbation states (Fig. 3a). For a single external stimulation, such as an electric shock or a single drug treatment, a learnable [PER] token represents the perturbation signal. For genetic perturbations, each gene is modeled as a learnable token, integrated through a Graph Neural Network (GNN) with Gene Ontology (GO)^47^ relationships as edges, enabling the prediction of unseen genetic perturbations (Fig. 3b).

**Fig. 3.**
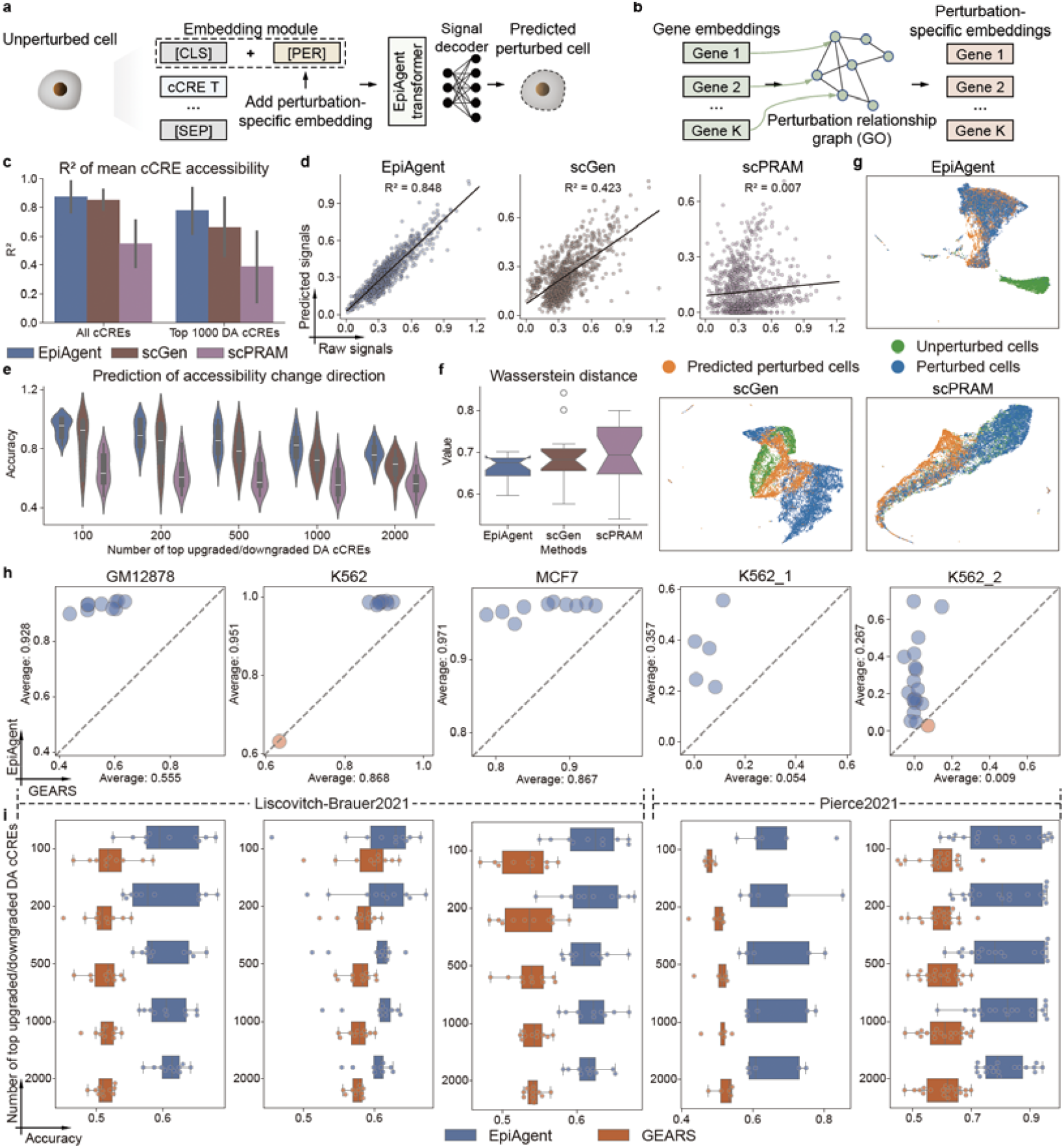
Prediction of cellular responses to out-of-sample stimulated and unseen genetic perturbations using EpiAgent. **a,** Workflow for predicting the post-perturbation state from the corresponding unperturbed cell using the fine-tuned EpiAgent model. **b**, Workflow for obtaining perturbation-specific embeddings for genetic perturbations. **c-g**, Benchmarking performance for prediction of cellular responses to out-of-sample stimulated perturbations. **c**, Coefficients of determination (R^2^) for mean cCRE accessibility between raw signals and predicted signals from different methods, evaluated across all cCREs and the top 1,000 DA cCREs. Bar heights denote the mean, and error bars denote the 95% confidence intervals. **d**, Linear regression fit for mean accessibility of the top 1,000 DA cCREs between raw signals and predicted signals from different methods, with mono-1 cells as query data. **e**, Prediction accuracy for accessibility change directions under varying numbers of top upgraded/downgraded DA cCREs. **f**, Wasserstein distance between raw signals and predicted signals from different methods for the top 1,000 DA cCREs. **g**, UMAP visualization of cell embeddings from different methods for unperturbed cells, perturbed cells, and predicted perturbed cells, with mono-1 cells as query data. **h & i**, Benchmarking performance for prediction of cellular responses to unseen genetic perturbations. **h**, Pearson correlation coefficients for mean cCRE accessibility between raw signals and predicted signals from different methods, evaluated for the top 1,000 DA cCREs. Each point represents the results for a specific gene perturbation. **i**, Prediction accuracy for accessibility change directions under varying numbers of top upgraded/downgraded DA cCREs. In **e & i**, the numbers on the axes indicate the sizes of the sets of upregulated and downregulated DA cCREs being evaluated. In **f & i**, the upper, middle, and lower edges of the boxes denote the upper quartile, median, and lower quartile, respectively, and error bars denote the maximum and minimum values, excluding outliers.

#### Out-of-sample stimulated perturbation

Lareau et al.^48^ used dsciATAC-seq to profile chromatin accessibility in untreated control and ex vivo cultured, lipopolysaccharide-stimulated bone marrow mononuclear cells (BMMCs). We conducted experiments in which perturbed cells of one cell type serve as the query dataset, and the remaining cells serve as the reference dataset. The process was repeated until all perturbed cells had been used as the query dataset. Notably, only the resting cells are in the Human-scATAC-Corpus, preventing any risk of data leakage. EpiAgent was fine-tuned on the reference dataset, and scGen^49^ and scPRAM^50^ were trained as baseline methods for comparison. Due to characteristics of high dimensionality and large cell count, other corresponding methods^51,52^ could not be executed successfully and were therefore excluded from the comparison. Details on experimental setup, fine-tuning of EpiAgent, and baseline methods are provided in Methods.

As shown in Fig. 3c, EpiAgent demonstrates its superior performance, showing high correlation between predicted and real signals of mean accessibility of all cCREs, surpassing scGen and scPRAM by an average of 2.504% and 59.240% for all cell types, respectively. For the top 1,000 differentially accessible (DA) cCREs, the advantages increase to 17.112% and 99.680%, highlighting the ability of EpiAgent to accurately predict accessibility changes specific and sensitive to perturbations. For example, in “mono-1” cells (Fig. 3d), average accessibility of DA cCREs predicted by EpiAgent closely aligns with the real signal, forming an almost perfect linear relationship, while scGen and scPRAM fail to capture this intrinsic pattern. Such a trend is also observed across other cell types (Supplementary Fig. 13). EpiAgent also shows remarkable accuracy in predicting the direction of accessibility changes for DA cCREs under perturbation. When DA cCREs are divided into upregulated and downregulated groups, EpiAgent consistently outperforms baselines in predicting the overall accuracy for varying numbers of top DA cCREs. For the most perturbation-sensitive cCREs, such as the top 100 upgraded/downgraded DA cCREs, EpiAgent achieves an average accuracy even exceeding 90%.

Beyond predicting overall signals, EpiAgent effectively captures heterogeneity of individual cells in responses to perturbations. By calculating the Wasserstein distance of top 1,000 DA cCREs between perturbed cells and the predicted by each method, we found EpiAgent gets lower distance values than those of scGen and scPRAM, indicating closer alignment of cellular distributions between the predicted and real signals. Visualization of cell embeddings (Fig. 3g and Supplementary Fig. 14) reveals that, in latent space of EpiAgent, predicted post-perturbation cells nearly overlap with the real while maintaining clear separation from pre-perturbation cells. In contrast, baseline methods either completely separate or overly merge the three cell populations in their latent space.

#### Unseen genetic perturbation prediction

Liscovitch-Brauer et al.^53^ and Pierce et al.^54^ developed CRISPR-sciATAC and Spear-ATAC technologies, respectively, enabling simultaneous capture of chromatin accessibility profiles and CRISPR perturbations. These datasets include chromatin accessibility changes in GM12878, K562, and MCF7 cells before and after various gene knockouts. Following GEARS^55^, we extracted 25% of perturbation-related cells from each dataset as the query dataset and fine-tuned EpiAgent on the remaining cells. GEARS was trained as a baseline method for comparison. As shown in Fig. 3h, EpiAgent outperforms GEARS in predicting changes of mean cCRE accessibility under different genetic perturbations. In the Liscovitch-Brauer2021 dataset, EpiAgent achieves an average Pearson correlation coefficient 24.454% higher than GEARS, while in the Pierce2021 dataset, GEARS even fails to predict perturbation-induced signals effectively (Fig. 3h). Compared to EpiAgent, GEARS lacks both a transformer architecture and a pretraining strategy, limiting its ability to accurately capture information from regulatory network and predict genetic perturbations at the epigenomic level. Such the limitation is also evident in the accuracy of predicting accessibility change directions for top upgraded/downgraded DA cCREs, where GEARS performed only slightly better than random guessing, while EpiAgent demonstrates significantly higher accuracy (Fig. 3i).

Collectively, all the aforementioned results highlight the superiority of EpiAgent to accurately characterize cellular heterogeneity under both stimulated and genetic perturbations by leveraging its unique transformer-based structure, pretraining strategy, and integration of perturbation-specific tokens.

### EpiAgent facilitates reference data integration and query data mapping

As previously demonstrated, EpiAgent can extract cell embeddings that encapsulate cellular heterogeneity from a single dataset. However, in practical scenarios such as constructing cell atlases, cells are typically derived from multiple datasets, and batch effects, that is external factors such as differences in sequencing technologies or depths, will be introduced and result in variations in sequencing data for the same cell types across datasets^56^. For the correction of batch effects, EpiAgent incorporates a batch-specific token embedding to generate batch-invariant cell embeddings, and once integrated, these corrected cells can serve as reference data (Fig. 4a). When newly-sequencing data is introduced as query data, EpiAgent generates cell embeddings that can be directly mapped to the reference dataset using mutual nearest neighbor (MNN) relationships, enabling downstream tasks such as label transfer (Fig. 4b).

**Fig. 4.**
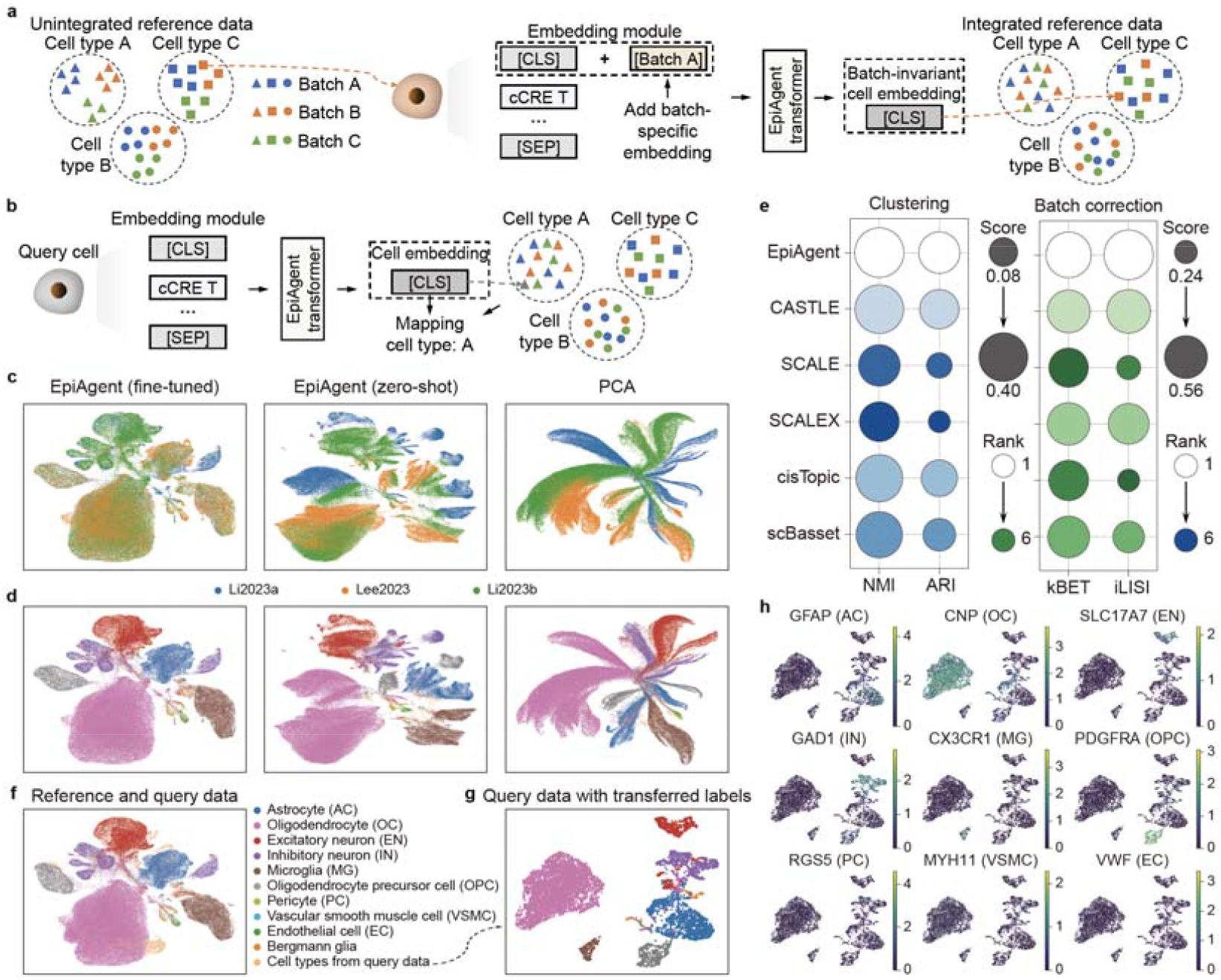
Reference data integration and query data mapping using EpiAgent. **a,** Workflow for the integration of reference data using the fine-tuned EpiAgent model. **b**, Workflow for mapping query data onto the integrated reference data using the fine-tuned EpiAgent model, followed by label transfer. **c-d**, UMAP visualization of the reference dataset using cell embeddings derived from the fine-tuned EpiAgent model, the zero-shot EpiAgent model, and PCA, labeled by batches (**c**) and cell types (**d**), respectively. **e**, Benchmarking performance of EpiAgent compared to baseline methods, evaluated using clustering metrics (NMI and ARI) and batch correction metrics (kBET and iLISI). **f**, Visualization of reference data and query data, with query cells colored separately. **g**, Visualization of query data with transferred labels. **h**, Marker gene expression for potential cell types in the query dataset. Cell embeddings for UMAP visualizations in panels **f-h** were derived from the fine-tuned EpiAgent model.

Here we constructed a reference dataset of the human brain using cells partially derived from the Li2023a^8^, Lee2023^57^, and Li2023b^39^ datasets, and notably, Li2023a and Lee2023 datasets are also part of the Human-scATAC-Corpus. Details on data filtering and preprocessing are provided in Methods. We fine-tuned EpiAgent on the reference dataset with the batch labels, and compared its performance to the zero-shot EpiAgent model and one of the most widely-used dimensionality reduction methods, PCA. As shown in Fig. 4c and 4d, without introducing batch information as labels, embeddings from the zero-shot EpiAgent model, similar to those derived from PCA, show pronounced batch effects between datasets, particularly between Li2023a and the other two datasets. In contrast, after fine-tuning, EpiAgent leverages its batch-specific token embedding and self-attention mechanism to successfully mitigate or even eliminate batch-specific differences in cells of the same type across datasets. We benchmarked the fine-tuned EpiAgent model against baseline methods on perspectives of clustering and batch correction, using NMI and ARI to assess clustering performance, and kBET and iLISI to evaluate batch correction performance. EpiAgent still achieves the best overall performance, demonstrating its ability to eliminate batch effects while preserving cellular heterogeneity (Fig. 4e, Supplementary Fig. 15 & 16). Among the baseline methods, CASTLE, SCALEX, and scBasset also include batch correction modules, enabling partial mitigation of batch effects, while SCALE and cisTopic lack the corresponding mechanisms, resulting in distinct batch-specific boundaries similar to those observed with PCA.

We further evaluated the query data mapping capability of EpiAgent using a single-cell multi-omics human brain datasets generated by 10X Genomics as query data (Fig. 4f). Query cells are mapped onto the reference dataset, with the labels of their 20 nearest neighbors in the reference dataset used for label transfer (Fig. 4g). The transferred labels were validated against the expression of marker genes for each cell type (Fig. 4h). Across nearly all cell types, the labels transferred by EpiAgent show strong concordance with marker gene expression. For major cell types, such as astrocytes, EpiAgent distinguishes them with high clarity; for rare cell types, such as endothelial cells, which contain very few cells and are difficult to be identified in single-dataset clustering, EpiAgent assigns them labels via mapping, surpassing the limitations of clustering within individual datasets. In summary, the fine-tuned EpiAgent model enables effective reference data integration by minimizing batch effects and accurate query data mapping without requiring additional training or batch-specific information, highlighting the potential as a robust tool for constructing large-scale, multi-dataset scATAC-seq cell atlases.

### EpiAgent enables in-silico treatment for cancers through key cCRE knockouts

Alterations in chromatin accessibility play a critical role in the initiation and progression of cancer, while identifying epigenetic drivers and quantitatively evaluating their effects on cancer remains a significant challenge^58,59^. Benefiting from the unique modeling for scATAC-seq data as cell sentences composed only of accessible cCRE indices, EpiAgent can directly perform the knockout of specific cCREs to observe resulting cellular changes. We extended the capability to the in-silico treatment for cancers, investigating whether key cCRE knockouts could potentially transform cancer cells towards a more normal-like state (Fig. 5a). For validation, we fine-tuned EpiAgent on a clear cell renal cell carcinoma (ccRCC) dataset, which reveals epigenetic regulatory processes between normal and cancer cells^60^. In their studies, Terekhanova et al.^60^ highlighted two regulatory regions associated with the ABCC1 and VEGFA genes that are active across multiple cancers, as well as a promoter specific to ccRCC corresponding to the EGLN3 gene. By visualizing the accessibility of all cCREs upstream of the transcription start sites (TSS) for these genes (ABCC1 and VEGFA: 2,000 bps upstream; EGLN3: 5,000 bps upstream), we identified their promoters, highlighted in red in Fig. 5b, as targets for knockout.

**Fig. 5.**
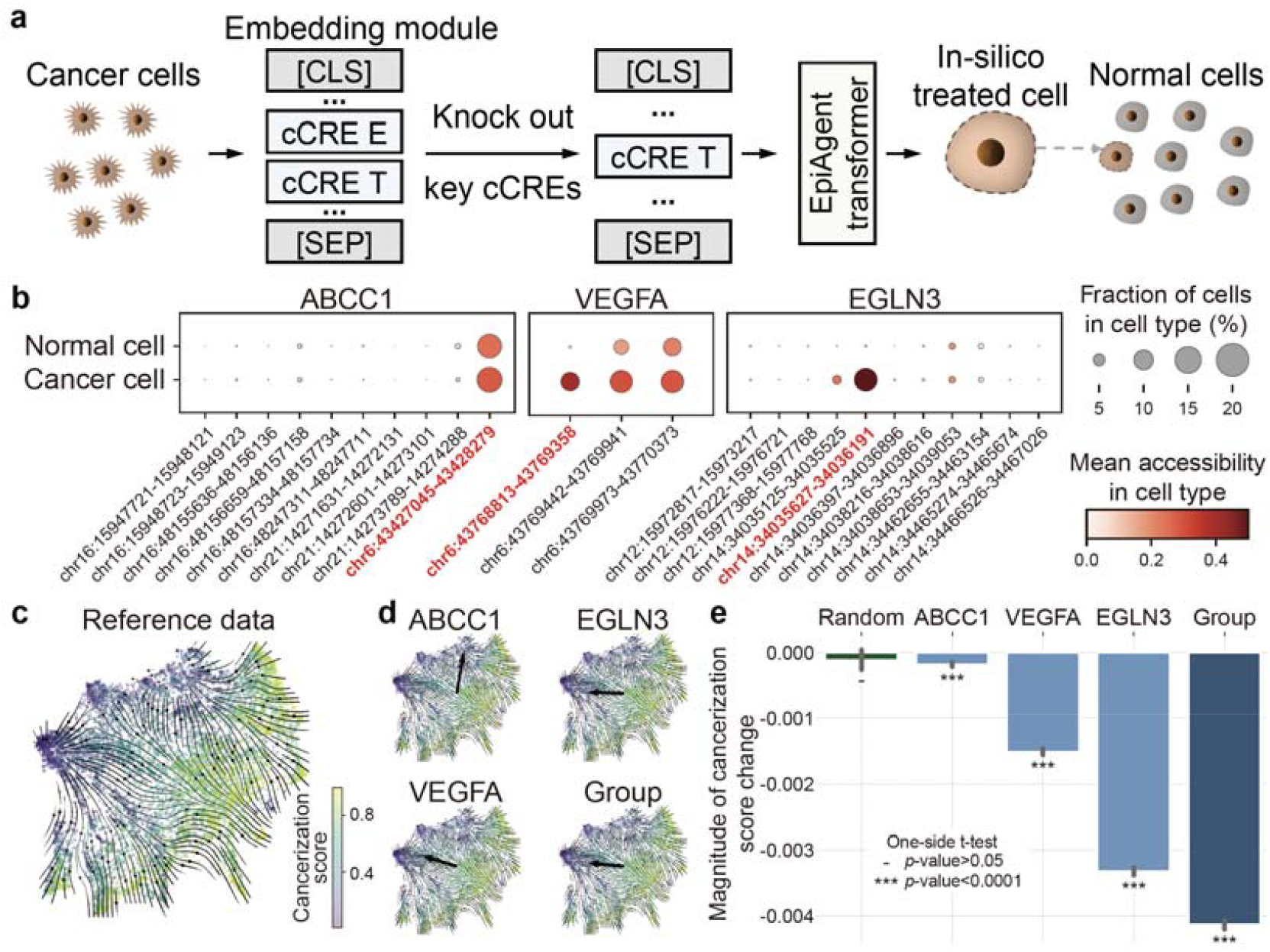
In-silico treatment for cancers through key cCRE knockouts using EpiAgent. **a,** Workflow for knocking out key cCREs using the fine-tuned EpiAgent model. **b**, Accessibility of cCREs upstream of the TSS for key genes in ccRCC (ABCC1 and VEGFA: 2,000 bp upstream; EGLN3: 5,000 bp upstream). Promoters identified as the target cCREs for knockout are highlighted in red. **c**, Visualization of the reference dataset comprising chimeric cells with varying cancerization scores. Manifold denote changes in cancerization scores, and arrows denote the direction from low to high cancerization levels. d, Directional shifts of query cells in response to knockouts of different promoters (arrows) against the background of the reference dataset. “Group” represents the combined knockout of the promoters for all three genes. **e**, Magnitude of cancerization score changes following individual promoter knockouts (ABCC1, VEGFA, EGLN3) and the combined knockout, compared to random cCRE knockouts. Bars represent mean values, and error bars denote the 95% confidence interval. One-sided t-tests are performed with the alternative hypothesis that the mean is less than 0.

To quantitatively evaluate the effects of knockout, we first constructed a synthetic dataset comprising chimeric cells with varying degrees of cancerization, serving as a reference. Specifically, for each chimeric cell, its cell sentence is generated by sampling and concatenating the cCRE indices from cell sentences of normal and cancer cells, and the proportion of cCREs from cancer cells is defined as its cancerization score (Fig. 5c). Upon the same procedure, we then generated a set of chimeric cells with a cancerization score of 0.5 as query cells, and for each query cell, we introduced the specific cCRE index into its cell sentence (if absent) and subsequently removed it to simulate the knockout process. Detailed steps of construction are provided in the Methods.

With the reference dataset as background, the results show that while knockout of the ABCC1 promoter resulted in an overall directional shift distinct from those induced by knockouts of other promoters individually or in combination (referred to as “Group”), all scenarios consistently shift cells from higher to lower cancerization scores (Fig. 5d). To quantify effects for all knock outs, we identified the nearest neighbors of each query cell in the reference dataset, both pre- and post-knockout, within the EpiAgent latent space, and the difference in the mean cancerization scores of these neighbors is the magnitude of cancerization score change that we defined. As shown in Fig. 5e, comparison against a control group (referred to as “Random”), in which random cCREs were knocked out, shows that knockouts of target promoters obtain significantly negative changes in cancerization scores (one-tailed t-test, *p*-value < 0.0001), while random knockouts result in changes centered around zero. Among these knockouts, the promoter of the EGLN3 gene, a ccRCC-specific key cCRE, demonstrates a far greater effect on reversing cancerization compared to cCREs broadly accessible across various cancers. Simultaneous knockout of these promoters results in a more pronounced reversal of cancerization compared to individual knockouts. The instance shows the ability of EpiAgent for in-silico treatment in ccRCC, and also highlights the potential for broader applications of EpiAgent in other cancers and diseases.

### EpiAgent-B and EpiAgent-NT directly annotate newly-sequencing datasets without additional training

The zero-shot capabilities of the pretrained EpiAgent model are limited to unsupervised feature extraction, relying on clustering and prior biological knowledge for cell type annotation. To enable direct annotation without additional training, we developed EpiAgent-B and EpiAgent-NT, tailored for cell annotation in brain tissues and non-brain tissues, respectively. Specifically, EpiAgent-B and EpiAgent-NT build on the pretrained EpiAgent model by replacing the signal decoder with a classifier for supervised learning tasks, trained on the Li2023a^8^ and Zhang2021^7^ datasets, respectively, using cell-type labels as supervision (Fig. 6a). To mitigate potential batch effects that could overshadow true cell type differences, we coarsened the granularity of the reference dataset labels. Detailed procedures for training and prediction of EpiAgent-B and EpiAgent-NT are provided in Methods.

**Fig. 6.**
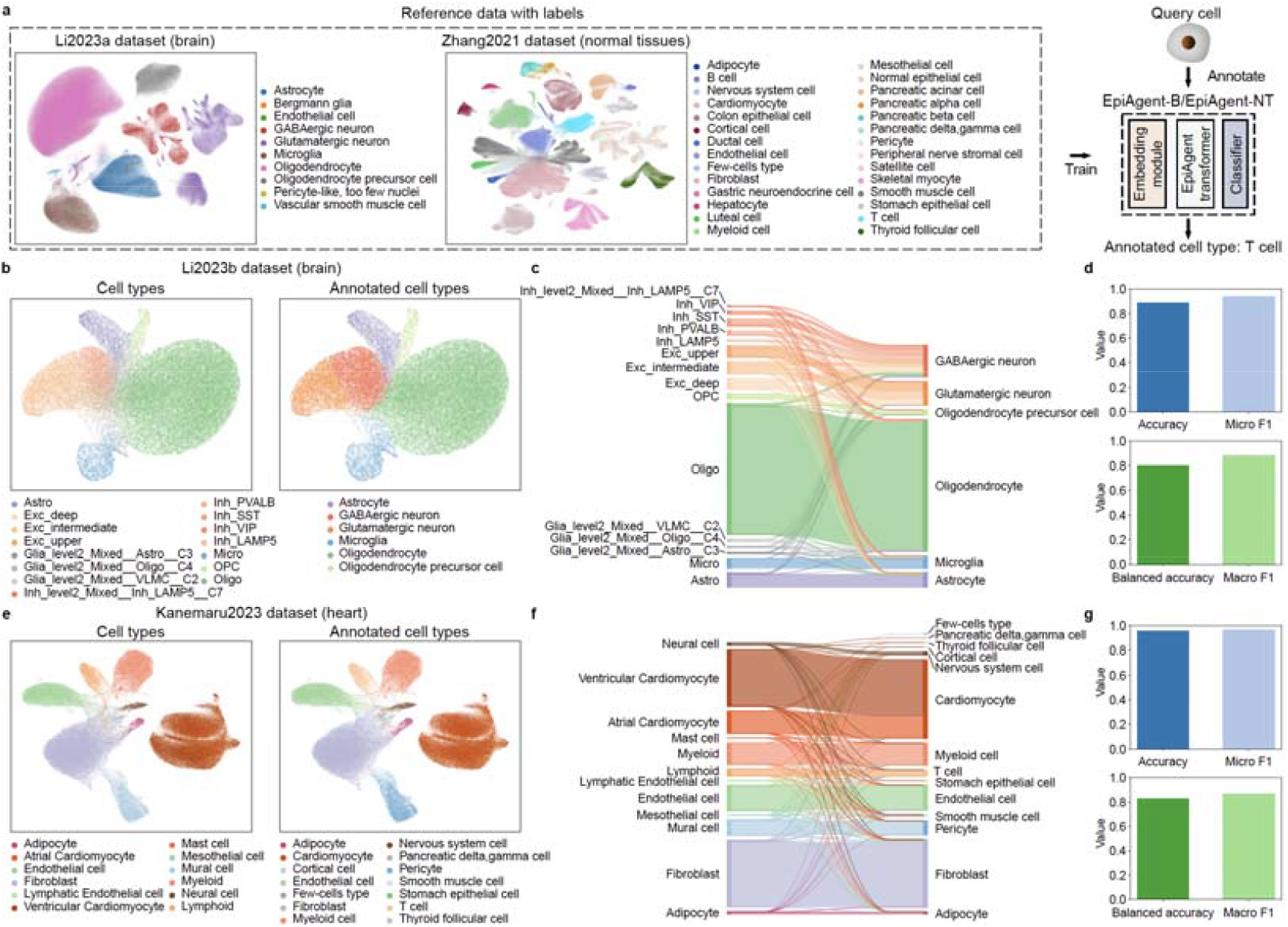
direct annotation on newly-sequencing datasets using supervised EpiAgent-B and EpiAgent-NT models. **a,** Overview of reference datasets used for training and the architecture of EpiAgent-B and EpiAgent-NT. The Li2023a and Zhang2021 datasets were used to train EpiAgent-B and EpiAgent-NT, respectively. Cell embeddings for UMAP visualization are derived from the pretrained EpiAgent model. **b-d**, Cell type annotation on the Li2023b dataset using EpiAgent-B. **e-g**, Cell type annotation on the Kanemaru2023 dataset using EpiAgent-NT. **b & e**, Visualization of the Li2023b (**b**) and Kanemaru2023 (**e**) datasets, comparing ground-truth labels (left) with predicted labels (right) generated by EpiAgent-B (**b**) and EpiAgent-NT (**e**). Cell embeddings for UMAP visualization are derived from EpiAgent-B (**b**) and EpiAgent-NT(**e**), respectively. **c & f**, Sankey plot for illustrating annotated results on the Li2023b (**c**) and Kanemaru2023 (**f**) datasets for each cell type. Ground truth labels are shown on the left, and predicted cell types using EpiAgent-B (**c**) and EpiAgent-NT (**f**) are shown on the right. **d & g**, Annotation performance of EpiAgent-B (**d**) and EpiAgent-NT(**g**) on the Li2023b (**d**) and Kanemaru2023 (**g**) datasets evaluated by accuracy, micro F1, balanced accuracy, and macro F1.

Compared to other methods, adapting EpiAgent for direct annotation poses several distinct advantages: the EpiAgent architecture is inherently designed for training on large-scale datasets, exceeding 100,000 cells, which is currently infeasible for most scATAC-seq-based methods; EpiAgent has been pretrained on the massive, unlabeled Human-scATAC-Corpus, which significantly enhances the robustness for annotation across various tissues and sequencing technologies; EpiAgent uses a standardized set of cCREs covering the majority of datasets, mitigating errors introduced by inconsistent features between datasets and bypassing inaccuracies stemming from converting cCRE accessibility into gene activity scores; the unique modeling for scATAC-seq data in EpiAgent, which exclusively considers accessible cCREs, allows it to maximize attention on cell-type-specific signals without the need for cCRE filtering.

For evaluation, we tested the performance of EpiAgent-B and EpiAgent-NT on the Li2023b and Kanemaru2023 datasets, respectively, demonstrating their superior ability for cell type annotation (Fig. 6b-g). The visualization of cell embeddings (Fig. 6b & e) shows that, compared to the unsupervised EpiAgent model (Supplementary Fig. 2 & 3), the supervised EpiAgent-B and EpiAgent-NT models tends to group cells of the same major type into a unified cluster rather than into separate subclusters. Although the change may sacrifice some subpopulation heterogeneity in the latent space, it significantly enhances the robustness and accuracy of EpiAgent-B and EpiAgent-NT for cell type annotation (Fig. 6c & f).

In terms of overall annotation performance, EpiAgent-B achieves an accuracy and micro F1 score exceeding 0.89 and 0.94, respectively, while EpiAgent-NT exceeds 0.95 for both metrics (Fig. 6d & g). When focusing on each cell type individually, both models demonstrate balanced accuracy and macro F1 scores above 0.8, indicating strong performance for both major and rare cell types. All the results highlight the effectiveness of EpiAgent-B and EpiAgent-NT for pre-annotating scATAC-seq datasets, suggesting that suggesting their potential to be valuable tools for scATAC-seq data pre-analysis.

## Discussion

Here, we introduce EpiAgent, the first foundation model specifically designed for scATAC-seq data. EpiAgent overcomes several key limitations of existing methods, including limited framework generalizability, underutilization of large-scale data, and inability to capture all accessible cCREs. By ranking accessible cCREs in each cell according to their importance, EpiAgent constructs cell sentences as input, thereby increasing information density while retaining critical regulatory signals. EpiAgent includes 1.4 billion parameters overall, with its core module (the EpiAgent transformer) containing approximately 56 million parameters. We pretrained EpiAgent on the Human-scATAC-Corpus, containing approximately 5 million cells and 35 billion tokens, using two newly designed tasks—the cell-cCRE alignment task and the signal reconstruction task—that are well-suited to data lacking explicit contextual relationships, allowing EpiAgent to capture both cellular heterogeneity and regulatory networks.

To validate the effectiveness, robustness, and versatility of EpiAgent, we conducted comprehensive benchmarking across multiple downstream tasks. In unsupervised feature extraction, the fine-tuned EpiAgent can generate cell embeddings that effectively capture cellular heterogeneity, and on datasets with cell populations resembling those in the Human-scATAC-Corpus, EpiAgent can achieve performance on par with or even surpassing baseline methods under zero-shot settings. For cell type annotation, EpiAgent outperforms baseline methods by enabling stable training on large-scale data and achieving consistent performance across diverse datasets. Leveraging the signal decoder, EpiAgent not only can reconstruct raw signals from cell embeddings, but also can filter noise and provide stable data imputation.

We further demonstrated that EpiAgent can incorporate additional information at the [CLS] token embedding to accurately predict cellular responses to single external perturbations. By substituting the perturbation-specific token embedding with the output from a GO-based graph neural network, EpiAgent can accurately predict cellular responses to unseen genetic perturbations. Similarly, by introducing batch-specific information into the [CLS] token embedding, EpiAgent can integrate multiple human brain datasets as reference data, enabling query-to-reference mapping for label transfer. Leveraging the cell-sentence framework, EpiAgent can also simulate in-silico cCRE knockouts in cancer cells, quantifying the impact of promoters of three cancer-related genes (ABCC1, VEGFA, and EGLN3) in ccRCC, thereby showcasing its potential utility for in-silico treatment. Additionally, we developed EpiAgent-B and EpiAgent-NT, models that can annotate cell types in newly sequenced data without additional training.

Moving forward, we also describe several directions for improving EpiAgent. First, by incorporating DNA sequences, we can unify cCREs from multiple species into a shared embedding space, enabling the development of a multi-species foundation model beyond human cells. Second, by integrating transcriptomic and other multi-omic data into the cell sentences, we can develop a comprehensive multi-omic foundation model capable of dissecting regulatory networks from multiple perspectives. Finally, by combining EpiAgent with foundation models from other domains, such as natural language processing and computer vision, we would pave the way for multimodal intelligent agents for cells, transcending the boundaries of omic data alone.

## Methods

### Data collection and preprocessing of Human-scATAC-Corpus

We manually curated 27 datasets from published literatures^6,7,9,39,48,57,61–78^, along with 10 additional PBMC dataset from the 10X Genomics website (https://www.10xgenomics.com/datasets/). Of these, 16 datasets include fragment data, while the remaining datasets only provide cell-by-peak matrices. Notably, since the data collected in the Human-scATAC-Corpus is unlabeled, conducting peak calling across all datasets could bias the identified cCREs towards cell types or tissues with larger cell numbers. Therefore, we did not perform peak calling directly on all datasets but instead merged publicly available cCRE sets. The curated cCRE set is derived from the following sources: an atlas-level dataset containing various human tissues (excluding adult brain)^7^, an atlas-level dataset from human brain tissues^8^, and a large-scale human blood and bone marrow dataset^48^. Overlapping cCREs were combined into longer intervals, and those accessible in fewer than 100 cells were excluded. The curated cCRE set contains 1,355,445 cCREs, and overlap analysis compared to the peak or cCRE sets from the published datasets demonstrates a high degree of concordance, indicating that our curation encompasses the majority of potential cCREs (Supplementary Fig. 17 & 18).

To filter out low-quality cells in the Human-scATAC-Corpus, we removed cells with fewer than 100 accessible cCREs. Both fragment data and cell-by-peak matrices were mapped to the predefined cCRE set. The data were then segmented into chunks of 5,000 cells each for pretraining purposes. Ultimately, the Human-scATAC-Corpus comprise a total of 5,000,000 cells across 31 tissues, with a total of 10 billion tokens (accessible cCREs), providing a comprehensive resource for pretraining EpiAgent. Detailed information on each preprocessed dataset is available in Supplementary Table 1.

### Non-zero ranked value tokenization

All scATAC-seq data in the Human-scATAC-Corpus are preprocessed into a large-scale cell-by-cCRE count matrix ***X*** ∈ ℝ^*N*×*P*^, where *N* and *P* denote the number of cells and the number of cCREs, respectively. Given that the raw count matrix ***X*** is quasi-binary, with the majority of elements being 0, 1, or 2, we applied a TF-IDF transformation to ***X***, aiming to measure the importance of each cCRE for a given cell based on its occurrence within both the individual cell and the entire Human-scATAC-Corpus. Assuming that ***X***_*ij*_ is the element of cCRE *j* in cell *i* in the matrix ***X***, the TF-IDF transformation can be formulated as:

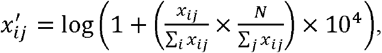

where 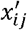 denotes the transformed element corresponding to ***X***_*ij*_ in the newly formed matrix ***X′*** Notably, the TF-IDF transformation does not alter the non-zero values in the cell-by-cCRE matrix to zero, nor does it transform zero values into non-zero values. Since the cell-by-cCRE matrix is extremely sparse and high-dimensional, during tokenization, we only consider cCREs with non-zero values. The ranks of these cCREs are determined by sorting them in the transformed cell vector 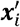 in descending order. Through the tokenization, each quasi-binary vector ***x***_*i*_·of an individual cell is converted into a cell sentence, that is a sequence of cCRE indices *C*_*i*_ = { *C*_*i*,1_, *C*_*i*,2…_}.

Since EpiAgent is a transformer-based model, the computational cost during forward and backward propagation is proportional to the square of the length of the cell sentence. To balance computational cost and the need to include as many tokens as possible per cell, we set the maximum tokenized cell sentence length to 8190 (excluding the [CLS] and [SEP] tokens). During each forward step, if the size of the cCRE index set exceeds the limit, random sampling without replacement is performed until the length constraint is met.

### Model architecture of EpiAgent

EpiAgent consists of three modules: an embedding module, the EpiAgent transformer, and a signal decoder. The embedding module converts the sequence of cCRE indices into a sequence of embeddings. The EpiAgent transformer takes these embeddings as input, and then generates contextual cCRE embeddings via a series of stacked transformer blocks. The output embedding corresponding to the [CLS] token serves as the cell embedding. The signal decoder takes the cell embedding as input, and reconstructs the raw signals of cCRE accessibility via a single-layer fully connected neural network.

### Embedding module

Embedding module consists of two main components: the cCRE embedding layer and the rank embedding layer. Before feeding the sequence of cCRE indices after tokenization into the model, two special tokens, [CLS] and [SEP], are added at the start and end, respectively, transforming the sequence into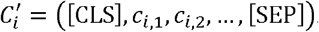. The cCRE embedding layer takes the processed sequence corresponding to each cell as input, and the rank embedding layer takes the sequence of rank indices *R*_*i*_ = (1,2,3,…) as input, which is a sequence of natural numbers with the same length as the cCRE index sequence for each cell. Embedding module then sums the outputs from the two embedding layers, which can be formulated as:

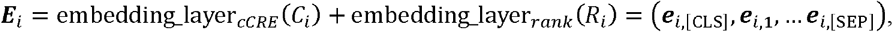

where embedding_layer_*cCRE*_(·) and embedding_layer _*rank*_ (·) denote the cCRE embedding and rank embedding layers, respectively.

Comparing the similar processing step to models in the NLP domain, the cCRE embedding and rank embedding layers are analogous to the word embedding and positional embedding layers, respectively. The key distinction is that the number of cCREs is significantly larger than the vocabulary size in NLP models, and the ranks represent the relative importance of each cCRE, derived from the values after the TF-IDF transformation, instead of raw positional locations in each cell. In embedding module, the dimensionality of embedding is set to 512. The cCRE embedding layer contains 1,355,449 embeddings, corresponding to 1,355,445 cCRE tokens and 4 special tokens. The rank embedding layer consists of 8,192 embeddings, which reflects the maximum token capacity (after adding the [CLS] and [SEP] tokens) of the EpiAgent transformer. The embedding module contains approximately 695 million parameters.

### EpiAgent transformer

As the core module of the transformer-based foundational model EpiAgent, the EpiAgent transformer consists of 18 stacked transformer blocks with a bidirectional attention mechanism to model global dependencies in high-dimensional input data, enabling the capture of epigenetic regulatory networks across different cells. Each transformer block operates on inputs and outputs with an embedding dimension of 512 and utilizes 8 attention heads. The input and output of each stacked transformer block can be formulated as:

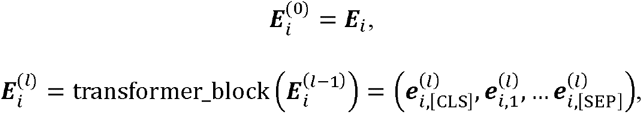

where 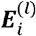 is the output of the *l*-th transformer block, and transformer_block (·) is a single transformer block. The core function of the transformer is to capture interactions between cCREs via the bidirectional attention mechanism, which can be formulated as follows:

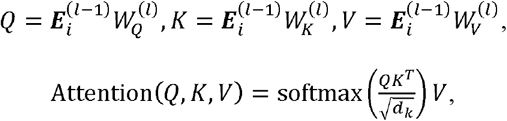

where 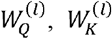 and 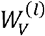 are trainable weight matrices used to compute the Query (Q), Key (K), and Value (V), respectively, and *d*_*k*_ denotes the dimension of the Key vector. For enhancing the representation capability for transformers, each transformer block leverages a multi-head attention mechanism by calculating multiple attention heads in parallel. Each attention head has its own set of 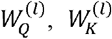 and 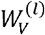, following the same formulation as single-head attention. The outputs from all attention heads are concatenated and then linearly transformed using a learnable matrix 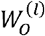, obtaining the output embeddings. The output of the final transformer block is 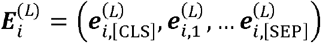, where 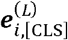 serves as the cell embedding of cell. The EpiAgent transformer contains approximately 57 million parameters and is implemented using the Flash Attention v2 framework to ensure efficient training and inference. Detailed parameters of the EpiAgent transformer are available in Supplementary Table 2.

### Signal decoder

Signal decoder employs a single-layer, fully connected neural network to transform cell embeddings into the accessibility signals of all cCREs for each cell. The process can be formulated as:

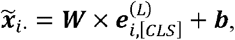

where ***W*** and ***b*** are learnable parameters of the single-layer neural network, and 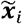·denotes the predicted accessibility signal for cell *i*, based on its cell embedding. The signal decoder contains approximately 695 million parameters.

### Pretraining tasks used for EpiAgent

To simultaneously capture gene regulatory networks and generate cell embedding rich in cellular heterogeneity, EpiAgent primarily utilizes two newly-designed tasks for pretraining: the cell-cCRE alignment task and the signal reconstruction task.

In the cell-cCRE alignment task, the aim is to train EpiAgent to identify potentially accessible cCREs that align with the regulatory networks compressed into the cell embedding. For cell *i*, we first obtain its cell embedding 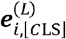 from EpiAgent. Assuming the length of the tokenized cell sentence *C*_*i*_ = (*C*_*i*,1_, *C*_*i*,2_,…) for cell *i* is *M*, we perform non-replacement sampling from cCREs that are inaccessible in cell *i* to construct a cCRE index set *B*_*i*_ = {*b*_*i*,1_, *b*_*i*,2_,…}, with a size of min (*α*_*alignment*_ *M*,*P* − *M*), where *α*_*alignment*_ is a pre-defined hyperparameter. We then input both *C*_*i*_ and *B*_*i*_ into the cCRE embedding layer, obtaining the input cCRE embeddings corresponding to the indices in these sets, denoted as ***É***_*i*_ = {***è***_*i*,1_, ***è***_*i*,2_,…} and ***É***_*i*_ **=** {***è***_*i*,1_, ***è***_*i*,1_,…}, respectively. Note that these embeddings are pre-transformer embeddings and are not influenced by the attention mechanism from other accessible or inaccessible cCREs within cell *i*, and thus only reflect information about the cCREs themselves. We then concatenate the cCRE embeddings from ***É***_*i*_ and ***É***_*i*_ with the cell embedding 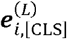, and merge these sets to form a new set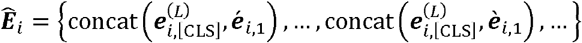. Each vector in this new set is then passed through a binary classifier to determine whether the cCREs in ***É***_*i*_ and ***É***_*i*_ are accessible in cell *i*, i.e., whether cell *i* aligns with the cCREs in ***É***_*i*_ and ***É***_*i*_. The task not only trains the self-attention mechanism but also engages more parameters from the embedding module, promoting faster convergence. The cell-cCRE alignment loss is computed using the “BCEWithLogitsLoss()” function in PyTorch, with the parameter “pos_weight” set to *α*_*alignment*_.

EpiAgent is pretrained for 12 epochs. During the first 4 epochs, we set *α*_*alignment*_ to 1; during the middle 4 epochs, we set *α*_*alignment*_ to 2; and for the final 4 epochs, we set *α*_*alignment*_ to 5. As the pretraining progresses, more cCREs are aligned with each cell.

In the signal reconstruction task, the aim is to enable the model to attend to all cCREs and recover the raw accessibility signals from the cell sentence. We first use the tokenized cell sentence *C*_*i*_ = (*C*_*i*,1_, *C*_*i*,2_,…)for cell *i* to generate a binary vector 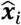, representing whether each cCRE in cell *i* is accessible. The predicted signal is the output 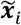 ·from the signal decoder, and 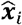·serves as the ground truth. The signal reconstruction loss is computed using “BCEWithLogitsLoss()” function in PyTorch, with the parameter “pos_weight” set to 100. The use of binary classification instead of regression for reconstruction task serves two purposes: first, it accelerates model convergence, and second, it conserves storage resources, as the Human-scATAC-Corpus only stores the cCRE indices for each cell, making classification tasks more storage-efficient.

Additionally, we observed that using only the two aforementioned tasks at the start of training could cause the pretraining to collapse. A potential reason is that these tasks are overly challenging during the warm-up phase of EpiAgent. To mitigate this, we additionally incorporated a replaced language modeling task during the first two epochs. Specifically, after tokenization, 50% of the cCREs (i.e., tokens) in each cell are randomly replaced by inaccessible cCREs, and the model then predicts whether each cCRE is accessible in the cell.

Although the EpiAgent transformer adopts an encoder architecture, it is not suitable for contextual pretraining tasks such as masked language modeling, primarily for three reasons. First, even after applying TF-IDF transformations to convert the quasi-binary cell-by-cCRE matrix into that with continuously-valued elements, many cCREs still remain inaccessible in each cell. In this context, whether a cCRE is accessible is more informative than subtle quantitative differences in its importance score. Second, the cCRE vocabulary is much larger than typical natural language vocabularies, making large-scale multi-class classification slow to converge and prone to collapse. Third, we aim for EpiAgent to not only leverage attention mechanisms to learn co-open patterns of cCREs, i.e., regulatory networks, but also to extract cell embeddings that capture cellular heterogeneity. Further details on the pretraining parameters can be found in Supplementary Table 2.

### Downstream analysis

#### Unsupervised feature extraction

In a downstream dataset, EpiAgent is fine-tuned using the same tasks as in pretraining, that is the cell-cCRE alignment task and the signal reconstruction task. The only difference in fine-tuning is the parameter *α*_*alignment*_ fixed to 1 for the cell-cCRE alignment task.

EpiAgent was compared against five baseline methods: cisTopic^14^, SCALE^15^, scBasset^17^, SCALEX^16^, and CASTLE^18^. All five baseline methods are well-designed and capable of processing large-scale datasets. We implemented each method according to their publications and GitHub documentation, ensuring that all parameters are consistent with the implementation in the publications or the default settings. For evaluation, we performed Leiden clustering on the cell embeddings obtained from each method, using a binary search approach to ensure that the number of clusters matches the actual number of cell types. Clustering performance is assessed using NMI and ARI scores. The detailed formulas for computing these metrics are available in Supplementary Note 2.

#### Supervised cell annotation

In the supervised cell annotation task, EpiAgent is fine-tuned by feeding the output cell embeddings into a classifier to perform multi-class classification. The classifier is a single-layer, fully connected neural network, and the loss function for fine-tuning is cross-entropy loss.

EpiAgent was compared against five baseline methods: PCA+SVM^42^, MLP^43^, EpiAnno^19^, CellCano^20^, and SANGO^21^. EpiAnno could not be executed on large-scale datasets using the parameters outlined in their publication. The limitation arises because EpiAnno does not account for batch effects and requires loading the entire dataset into the probabilistic graph during training. CellCano also encounters memory issues with large datasets. To address these constraints, we downsample cells when running both methods on large-scale datasets, keeping all other parameters unchanged. Compared to feature downsampling, cell downsampling leads to better performance for both EpiAnno and CellCano (Supplementary Fig. 19). Since PCA+SVM and MLP are methods specifically developed for scRNA-seq data, we convert the scATAC-seq data into gene activity matrices as inputs, aligning with the input used for Cellcano. Other parameters are set to be consistent with the implementations in their publications or the default settings. For each dataset, two-thirds of the cells are randomly assigned to the reference dataset, while the remaining cells serve as the query dataset. All methods are trained on the reference dataset and evaluated on the query dataset. We use accuracy and macro F1 for evaluation metrics, with the detailed formulas provided in Supplementary Note 2.

#### Data imputation

For the data imputation task, the fine-tuning strategy of EpiAgent is also consistent with the tasks used during pretraining, that is the cell-cCRE alignment task and the signal reconstruction task. The differences in fine-tuning are that, for the cell-cCRE alignment task, *α*_*alignment*_ is set to 1, and for the signal reconstruction task, mean squared error (MSE) loss is used for regressing the real signal.

EpiAgent was compared against two baseline methods: scCASE^23^ and scOpen^22^. First, we perform data imputation with each method and then apply PCA to the imputed data to obtain cell embeddings. Next, we conduct Leiden clustering on these embeddings and use a binary search procedure to ensure that the number of resulting clusters matches the true number of cell types. Clustering performance is assessed using the NMI and ARI scores, where better clustering results indicate superior performance on data imputation. The detailed formulas for computing these metrics are available in Supplementary Note 2. To make a fair comparison, we standardize the size of the cell-by-cCRE matrices used for data imputation across all methods. Specifically, we use a unified data preprocessing pipeline, retaining only cCREs accessible in at least 3% of cells. When processing datasets with more than 50,000 cells, scCASE triggers a “Segmentation fault (core dumped)” error, and scOpen encounters memory issues. In these cases, we randomly downsample to 50,000 cells and benchmark all methods on this reduced set. All other parameters are set in accordance with the implementations described in their publications or the default settings.

#### Out-of-sample stimulated perturbation prediction

In the task of out-of-sample stimulated perturbation prediction, the fine-tuning process for EpiAgent is as follows. First, we input all cells from the reference dataset into the pretrained EpiAgent model to obtain their cell embeddings, and employ Optimal Transport (OT) to match each perturbed cell with an unperturbed cell for each cell type. The detailed procedure for OT is available in Supplementary Note 3. Next, we construct a new reference dataset by duplicating data of unperturbed cells. For cells from the original dataset, we use the same training strategy as in data imputation (cell-cCRE alignment and signal reconstruction tasks) to allow EpiAgent to learn accessibility patterns of all cCREs. For the duplicated unperturbed cells, EpiAgent is trained to predict the post-perturbation states. Specifically, before feeding the input into the EpiAgent transformer, we introduce a learnable perturbation-specific token ([PER]) embedding in addition to the [CLS] token embedding. After obtaining cell embeddings through the EpiAgent transformer, the model performs cell-cCRE alignment and signal reconstruction tasks using the matched post-perturbation data as ground truth. For reconstructing original signals, the loss used for these two tasks is same as that in data imputation. To prevent the model from conflating unperturbed and perturbed data, we introduce a binary classification task during fine-tuning: one class consists of the original unperturbed cells, while the other class includes both the perturbed cells and the duplicated unperturbed cells. Under the training strategy, EpiAgent could leverage the perturbation-specific token and the attention mechanism to capture perturbation-related information. For unperturbed cells in the query dataset, EpiAgent can use their tokenized cell sentences (the sequences of cCRE indices) as input, along with the perturbation-specific token, to generate the predicted accessibility profiles of the corresponding perturbed cells.

We compare the performance of EpiAgent with two baseline methods: scGen^49^ and scPRAM^50^. We also considered including CellOT^79^ and scPreGAN^52^ as baseline methods. However, CellOT lacks GPU acceleration and failed to converge within a week on the dataset that we use for evaluation, and scPreGAN encountered a GPU memory issue due to its requirement to load the entire dataset into GPU memory at once. Given that scGen and scPRAM are designed for scRNA-seq data, which are not well-suited for processing the high-dimensional scATAC-seq data in its entirety, we standardize the dataset preprocessing, retaining only the top 50,000 most accessible cCREs as input for all methods. For evaluation, data from one cell type is selected as the query dataset, while data from other cell types are used as the reference dataset for training, repeating until all cell types are used as query datasets. We adopted three main evaluation criteria for benchmarking methods. First, to assess the prediction of average post-perturbation accessibility patterns, we computed the mean accessibility signals for both the real and predicted perturbed cells across all cCREs and the top 1,000 cCREs. Specifically, we fit a linear regression model to these mean values and used the resulting R^2^ as a performance metric. Second, to evaluate predictions of directional changes in accessibility, we identified an equal number of top up-graded and top down-graded DA cCREs from the query dataset, and then measured the accuracy of each method in capturing the correct direction of these accessibility changes. Third, to evaluate the prediction of cellular heterogeneity for perturbed cells, we used the Wasserstein distance between the predicted and real perturbed cells over the 1,000 DA cCREs, as a metric. Wasserstein distance is calculated by Euclidean distance based on cell-to-cell matching by OT, with the lower value indicating the better performance. The DA cCREs were determined with the “tl.rank_features” function in the epiScanpy^12^ pipeline, based on comparisons of unperturbed and perturbed cells in the query dataset.

#### Unseen genetic perturbation prediction

For the unseen genetic perturbation prediction task, the fine-tuning process of EpiAgent proceeds as follows. Given a genetic perturbation set *G* = {*g*_1_,*g*_2,…_}, where each element in *G* represents a perturbation corresponding to the knockout of a specific gene. We input each cell from the reference dataset into the pretrained EpiAgent model to obtain their cell embeddings, and for each gene perturbation, we perform OT to match the perturbed cells with their unperturbed counterparts based on the cell embeddings. Our fine-tuning aim is to enable EpiAgent to predict the chromatin accessibility landscape of a cell under a given gene perturbation, conditioned on the accessibility profile of its unperturbed state and a perturbation-specific token embedding. To achieve this, we introduce an additional embedding layer of genetic perturbation and a GNN into EpiAgent. The embedding layer contains a set of learnable embeddings ***P*** = {***p***_**1**_,***p***_**2**_,…}, where each element in ***P*** is a 512-dimensional vector corresponding to a specific perturbation (gene) in *G*. These embeddings are then processed by the GNN, which incorporates pathway information from GO^47^ to establish gene-gene and thus perturbation-perturbation relationships.

During fine-tuning, EpiAgent adds the perturbation-specific embedding on the input [CLS] token embedding of the unperturbed cell, and predicts the signals of the matched perturbed cell using the same loss as in the data imputation. Unperturbed and perturbed cells also undergo the cell-cCRE alignment and signal reconstruction tasks independently, without incorporating the perturbation-related embedding. By integrating these new embedding layers and the GNN, the fine-tuned EpiAgent model can generate perturbation-related token embeddings for previously unseen genetic perturbations and predict their chromatin accessibility effects when presented with chromatin accessibility profile of an unperturbed cell.

We select GEARS^55^ as the baseline method for comparison. Since GEARS is designed for scRNA-seq data and cannot process the high dimensionality of full scATAC-seq data, we preprocessed the test datasets to retain only the top 50,000 most accessible cCREs. We adopted two main evaluation criteria for benchmarking methods. First, to assess the prediction of average post-perturbation accessibility patterns, we compute Pearson correlation coefficients between the average accessibility signals of the top 1,000 DA cCREs in the real and predicted perturbed cells. Second, we evaluated predictions of directional changes in accessibility, with the same procedure in out-of-sample stimulated perturbation prediction. The DA cCREs were determined with the “tl.rank_features” function in the epiScanpy pipeline, based on comparisons of unperturbed and perturbed cells in the query dataset.

#### Reference data integration and query data mapping

In the reference data integration task, the fine-tuning process for EpiAgent is as follows. We input all the data into the pretrained EpiAgent model to obtain cell embeddings, and then use OT to match cells across different datasets (also served as batches). The detailed procedure for OT is available in Supplementary Note 3. Since there may be differences in cell populations between datasets, we apply a filtering process to the matched cell pairs. For each dataset pair, we calculate the distance between each matched pair and then train a Gaussian Mixture Model (GMM) with two components to classify the pairs. We consider the pairs belonging to the Gaussian distribution with the smaller mean as high-quality matches and retain them, while discarding the rest. For each batch of data, before inputting them into the EpiAgent transformer, we add a batch-specific token (e.g., [Batch 1], [Batch 2], [Batch 3], etc.) embedding on the [CLS] token embedding. The batch-specific tokens are designed to capture non-biological variations, such as technical differences. During fine-tuning, EpiAgent performs both the cell-cCRE alignment and signal reconstruction tasks, while the ground truths for the two tasks are derived not only from the cells themselves but also from their matched counterparts in other batches. Cells that do not have a match in other batches only use their own data for the ground truth. When generating cell embeddings, EpiAgent can optionally include or exclude the batch-specific token to decide whether or not the resulting embeddings eliminate batch-specific differences. We use NMI and ARI as clustering metrics, and kBET^80^ and iLISI as batch correction metrics. Clustering is performed using Leiden clustering, with the number of clusters set to match the actual number of cell types. The calculation of NMI and ARI is detailed in Supplementary Note 2, and kBET and iLISI were computed directly using the original code from scIB^81^.

In the query data mapping task, we assume the reference data has already been constructed by the fine-tuned EpiAgent model. For the query data, we input it directly into the fine-tuned EpiAgent model without the batch-specific token to obtain cell embeddings for the query data. Using MNN, EpiAgent identifies the 20 nearest neighbors in the reference dataset for each query cell in the cell embedding space, thus mapping the query data to the reference data. Then we can transfer the labels from reference data to query data, by assigning the label of the majority among the 20 nearest reference cells to each query cell.

#### In-silico treatment

For in-silico treatment, the process for EpiAgent is as follows. We first fine-tune EpiAgent on the cancer dataset using the same tasks as in unsupervised feature extraction, obtaining embeddings for both cancer cells and normal cells. Using the cell embeddings, we then use Leiden clustering on these cells to remove outliers or misannotated cells, and perform OT to match each cancer cell with a corresponding normal cell. For each matched pair, we create a synthetic cell with its cell sentence derived from randomly sampling cCRE indices from both cancer and normal cells, and the proportion of cCREs from the cancer cell defines the cancerization score. We set cancerization scores uniformly distributed between 0 and 1 to construct a reference dataset, and these synthetic cells are input into the fine-tuned EpiAgent model to generate embeddings for quantitative analysis.

To simulate a cCRE knockout for a given target cCRE, we construct a query dataset of synthetic cells with a cancerization score fixed at 0.5. If a target cCRE does not appear in the sentence of a synthetic cell, we add it. We then remove the target cCRE from the cell sentence to simulate its knockout. We input the cells before and after knockout into the fine-tuned EpiAgent model to obtain their embeddings. We then identify the 20 nearest neighbors of each query cell from the reference dataset before and after knockout, and measure how average cancerization scores of neighbors change. The change quantifies the magnitude of the cancerization score change resulting from the cCRE knockout.

#### EpiAgent-B and EpiAgent-NT model

Building on the pretrained EpiAgent model, we trained two additional models—EpiAgent-B and EpiAgent-NT—designed to directly annotate query data without the need for collecting reference data for training. Specifically, EpiAgent-B is used to annotate cells from brain tissues, whereas EpiAgent-NT targets cells from normal tissues excluding the brain. The training data are sourced from two well-annotated datasets in the Human-scATAC-Corpus: the Li2023a^8^ dataset, a cell atlas of brain tissues, and the Zhang2021^7^ dataset, a cell atlas of human normal tissues excluding the brain.

Before training, we aggregated cell type labels from these two datasets, and the mapping relationships between cell types before and after aggregation are detailed in Supplementary Table 3 and 4. The preprocessing step is necessary because overly fine-grained cell annotations could lead to batch effects overshadowing true cell type differences between the query and training data, potentially causing misannotations in the query data. The settings for training and parameters for classifier are consistent with those used in the supervised cell annotation task. In addition to the accuracy and macro F1 metrics used in the supervised cell annotation task, we also introduced micro F1 and balanced accuracy to further evaluate overall annotation performance as well as performance across each cell type. Detailed formulas for the metrics are provided in Supplementary Note 2.

## Data availability

The Human-scATAC-Corpus includes 27 published datasets^6,7,9,39,48,57,61–78^ and 10 publicly available PBMC datasets from the 10x Genomics website (https://www.10xgenomics.com/datasets/). All these datasets are publicly accessible, and detailed statistics and data availability are provided in Supplementary Table 1. Additional datasets^37–40,48,53,54,60^ not included in the Human-scATAC-Corpus, but used for evaluating EpiAgent in downstream analyses, are also publicly available, with detailed statistics and data availability provided in Supplementary Table 5.

## Code availability

The EpiAgent is freely available on Github (https://github.com/xy-chen16/EpiAgent).

## Acknowledgements

This work was supported by the National Key Research and Development Program of China (grant nos. 2023YFF1204802 and 2021YFF1200902) and the National Natural Science Foundation of China (grant nos. 62273194).

## Author contributions

R.J. conceived the study and supervised the project. X.C. collected and processed all data in Human-scATAC-Corpus and downstream analyses, and designed, implemented and validated EpiAgent. K.L. assisted in analyzing the results for data imputation, prediction of unseen genetic perturbations, and in-silico treatment. X.C. contributed to analyzing the results for unsupervised feature extraction and reference data integration. Z.W. helped with analyzing the results for supervised cell type annotation and validation of EpiAgent-B and EpiAgent-NT. Q.J. aided in the analysis prediction of out-of-sample stimulated perturbations. J.L. contributed to designing the pretraining tasks. Z.L. and Z.G. helped with implementing EpiAgent. X.C., K.L. and R.J. wrote the manuscript, with input from all the authors.

