## Supplementary file for "EpiAgent: Foundation model for single-cell epigenomic data"

**Supplementary Notes**

**Supplementary Note 1: Cellular pseudotime inference using ClockDML and cell embeddings derived from EpiAgent**

In the study by Xiao et al., it was noted that a higher proportion of ClockDMLs within accessible chromatin regions of a cell corresponds to a lower mitotic age. To obtain the mitotic age, which can also be used as pseudotime, we first calculated the total number of ClockDMLs in accessible cCREs for each cell. Compared with the original peak/cCRE set used in the dataset, the cCRE set used by EpiAgent provides broader genomic coverage, enabling more accurate initial estimates. To account for sequencing depth, we then normalized the total accessible ClockDML count by the number of accessible cCREs per cell, thereby obtaining a normalized value of accessible ClockDMLs.

Since scATAC-seq data exhibit inherently low signal-to-noise ratios, and many accessible regions—whether containing ClockDMLs or not—remain undetected, directly using these normalized values to infer pseudotime would be unreliable. Therefore, we leveraged cell embeddings derived from EpiAgent, which effectively capture cellular heterogeneity, to address the issue. We first computed the cosine similarity between each pair of cells based on embeddings and constructed a graph by connecting cells with similarity scores above 0.9 (including self-links). Each edge was weighted by the corresponding similarity score. Using the graph, we derived a state transition matrix by normalizing edge weights so that the sum of edges from each cell equals one. We then iteratively applied this transition matrix to the initial normalized accessible ClockDML values until convergence, obtaining final results for each cell.

Finally, we scaled the resulting values into the [0, 1] interval, mapping the highest estimated accessible ClockDML value to 0 and the lowest to 1. The resulting scale defines the pseudotime of each cell.

**Supplementary Note 2: Detailed formulas for all the metrics**

We employed the Normalized Mutual Information (NMI) and the Adjusted Rand Index (ARI) to assess clustering performance. Let $\boldsymbol{U}$ represent the ground-truth labels and $\boldsymbol{V}$ the predicted cluster labels. The NMI score is computed as follows:

$$NMI=\frac{MI\left( \boldsymbol{U},\boldsymbol{V} \right)}{\sqrt{H\left( \boldsymbol{U} \right)\cdot H\left( \boldsymbol{V} \right)}}$$

where $\mathrm{MI}\left( \cdot,\cdot\right)$ denotes the mutual information between the two label sets, and $H\left( \cdot\right)$ denotes the entropy. To calculate the ARI, let $u_{i}$ be the $i$th true label, $v_{j}$ the $j$th predicted label, $n_{ij}$ the number of spots assigned to both $u_{i}$ and $v_{j}$, $n_{i\cdot}$ the number of spots assigned to $u_{i}$, $n_{\cdot j}$ the number of spots assigned to $v_{j}$, and $n$ the total number of spots. The ARI score is computed as follows:

$$ARI=\frac{\sum_{i,j} \left( \begin{aligned} n_{ij} \\ 2 \end{aligned} \right)-\left[ \sum_{i} \left( \begin{aligned} n_{i\cdot} \\ 2 \end{aligned} \right)\sum_{j} \left( \begin{aligned} n_{\cdot j} \\ 2 \end{aligned} \right) \right]/\left( \begin{aligned} n \\ 2 \end{aligned} \right)}{\frac{1}{2}\left[ \sum_{i} \left( \begin{aligned} n_{i\cdot} \\ 2 \end{aligned} \right)+\sum_{j} \left( \begin{aligned} n_{\cdot j} \\ 2 \end{aligned} \right) \right]-\left[ \sum_{i} \left( \begin{aligned} n_{i\cdot} \\ 2 \end{aligned} \right)\sum_{j} \left( \begin{aligned} n_{\cdot j} \\ 2 \end{aligned} \right) \right]/\left( \begin{aligned} n \\ 2 \end{aligned} \right)}$$

We evaluated cell type annotation performance using accuracy and macro F1. Let the total number of cells be $N$, and among them, let the number of correctly annotated cells be $N_{\mathrm{correct}}$. Accuracy is defined as:

$$\mathrm{Accuracy}=\frac{N_{\mathrm{correct}}}{N}$$

For macro F1, let there be $C$ cell types. For each cell type $c\in\left\{ 1,2,\ldots,C \right\}$, we first compute precision ($P_{c}$) and recall ($R_{c}$). The F1 score for cell type $c$ is:

$$F1_{c}=\frac{2P_{c}R_{c}}{P_{c}+R_{c}}$$

Then the macro F1 is the average of the cell type-specific F1 scores:

$$Macro F1= \frac{1}{C}\sum_{c=1}^{C} {F1}_{c}$$

To further validate the performance of EpiAgent-B and EpiAgent-NT, we employed micro F1 and balanced accuracy. Micro F1 aggregates the contributions of all cell types to compute a single F1 score. Let $\sum\mathrm{TP}$, $\sum FP$, and $\sum\mathrm{FN}$ denote the total true positives, false positives, and false negatives across all cell types, respectively. Micro precision ($P_{\mathrm{micro}}$) and micro recall ($R_{\mathrm{recall}}$) are computed as:

$$P_{\mathrm{micro}}=\frac{\sum\mathrm{TP}}{\sum\mathrm{TP}+\sum FP}, R_{\mathrm{recall}}=\frac{\sum\mathrm{TP}}{\sum\mathrm{TP}+\sum\mathrm{FN}}$$

Since micro precision equals micro recall, the micro F1 score is simply:

$$micro F1=\frac{2P_{\mathrm{micro}}R_{\mathrm{recall}}}{P_{\mathrm{micro}}+R_{\mathrm{recall}}}=P_{\mathrm{micro}}=R_{\mathrm{recall}}$$

Balanced accuracy addresses cell type imbalance by considering the average recall across cell types. For the case of multiple cell types, let $R_{c}$ denote the recall for cell type $c$, and let there be $C$ cell types:

$$\mathrm{Balanced}a\mathrm{ccuracy}=\frac{1}{C}\sum_{c=1}^{C} R_{c}$$

**Supplementary Note 3: Detailed procedure of Optimal Transport for cell mapping**

Optimal transport (OT) aims to identify an optimal mapping that minimizes the cost of transforming one probability distribution into another. In our setting, we consider two sets of cells, represented by their cell embeddings $\boldsymbol{Z}_{A}\in\mathbb{R}^{n\times d}$ and $\boldsymbol{Z}_{B}\in\mathbb{R}^{n\times d}$, where $n$ and $m$ are the number of cells in the two sets and $d$ is the dimension of cell embeddings. We assume that both sets are uniformly distributed:

$$\boldsymbol{a}=\left( \frac{1}{n},\frac{1}{n},\ldots,\frac{1}{n} \right)\in\mathbb{R}^{1\times n}, \boldsymbol{b}=\left( \frac{1}{m},\frac{1}{m},\ldots,\frac{1}{m} \right)\in\mathbb{R}^{1\times m}$$

We construct a cost matrix $\boldsymbol{C}\in\mathbb{R}^{n\times m}$ using the Euclidean distances between the cell embeddings of the two sets:

$$\boldsymbol{C}=\mathrm{EuclideanDistance}(\boldsymbol{Z}_{A}, \boldsymbol{Z}_{B}).$$

The goal is to find an optimal coupling matrix $\boldsymbol{M}$ that minimizes the total transport cost. Formally, the optimal transport problem is defined as:

$$\boldsymbol{M}=\underset{\boldsymbol{M}}{arg min} {<\boldsymbol{M},\boldsymbol{C}>}_{F}$$

subject to:

$$\boldsymbol{M}\boldsymbol{\cdot1=}\boldsymbol{a}, \boldsymbol{M}^{\boldsymbol{T}}\boldsymbol{\cdot1=}\boldsymbol{b}, \boldsymbol{M}\boldsymbol{\geq}\mathbf{0}$$

where ${<\boldsymbol{M},\boldsymbol{C}>}_{F}$ denotes the Frobenius inner product. The minimal transportation cost obtained from this formulation is known as the Wasserstein distance, widely used as a measure of similarity between distributions.

The problem can be solved using the Sinkhorn algorithm, which introduces an entropic regularization term. Letting $\boldsymbol{K}=e^{-\frac{C_{ij}}{\varepsilon}}$, the algorithm alternately updates $\boldsymbol{u}$ and $\boldsymbol{v}$ until convergence:

$$u^{\left( l+1 \right)}=\frac{\boldsymbol{a}}{\boldsymbol{K}\boldsymbol{v}^{(l)}}, v^{(l+1)}=\frac{\boldsymbol{b}}{\boldsymbol{K}\boldsymbol{u}^{(l)}}.$$

After convergence, we obtain $\boldsymbol{u}^{*}$ and $\boldsymbol{v}^{*}$, and the approximate optimal coupling matrix can be expressed as:

$$\boldsymbol{M}=\mathrm{Diag}(\boldsymbol{u}^{*})\boldsymbol{K}\mathrm{Diag}(\boldsymbol{v}^{*})$$

The element $m_{ij}$ in $\boldsymbol{M}$ represents the proportion of mass assigned from cell $i$ in the first set to cell $j$ in the second set. Larger values of $m_{ij}$ indicate a closer match between the corresponding cell embeddings. For each row of $\boldsymbol{M}$, we select the column index with the maximum value, designating the paired cell in the second set that best aligns with the corresponding cell in the first set.

We implemented the OT-based cell matching using the Python Optimal Transport (POT) library. We applied the Euclidean metric and set numItermax=20000000 to ensure convergence and accuracy in the computation of the optimal coupling matrix.

**Supplementary Figures**

**Supplementary Figure 1**

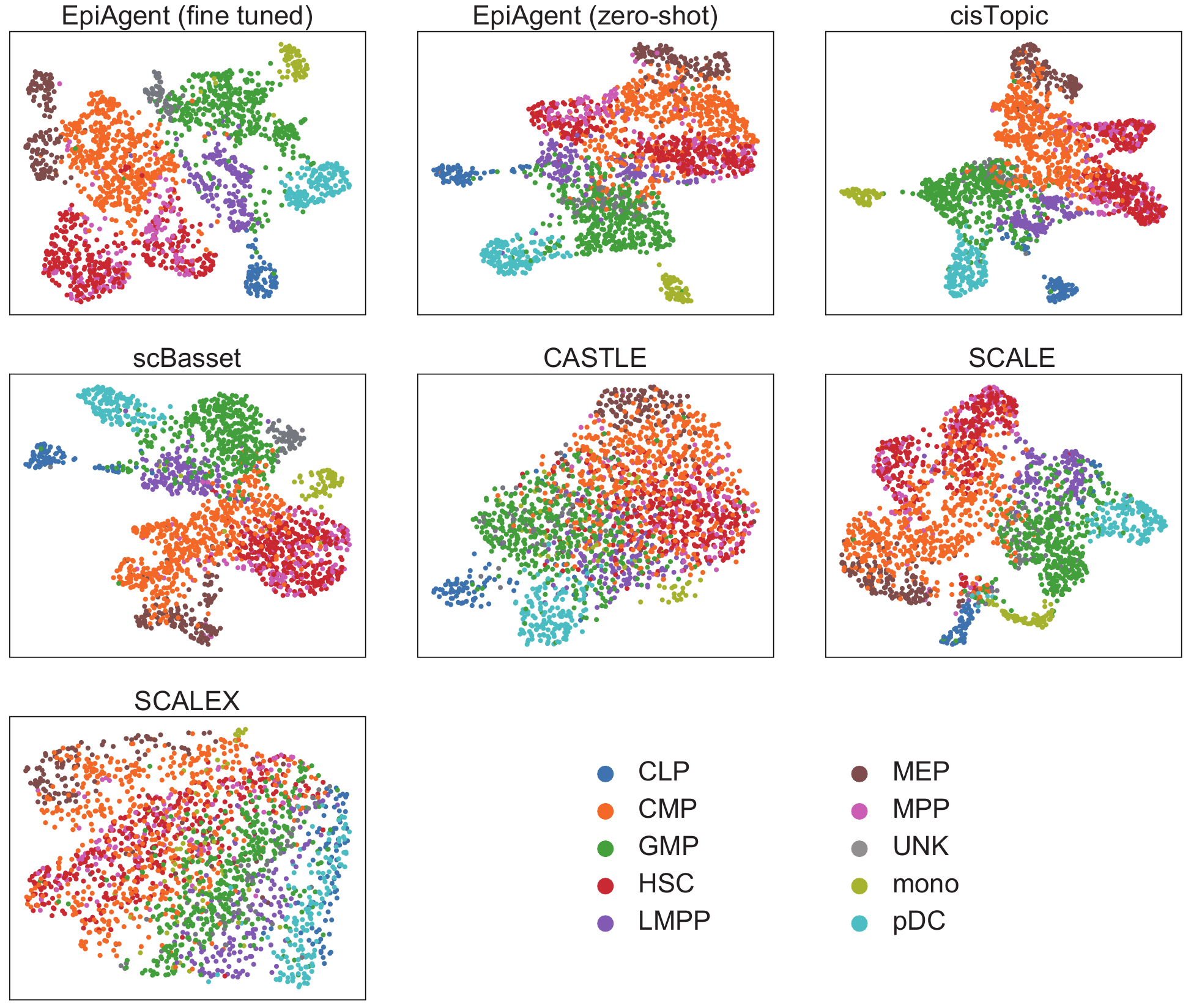

**Supplementary Figure 1.** UMAP visualization of cell embeddings derived from different methods on the Buenrostro2018 dataset.

**Supplementary Figure 2**

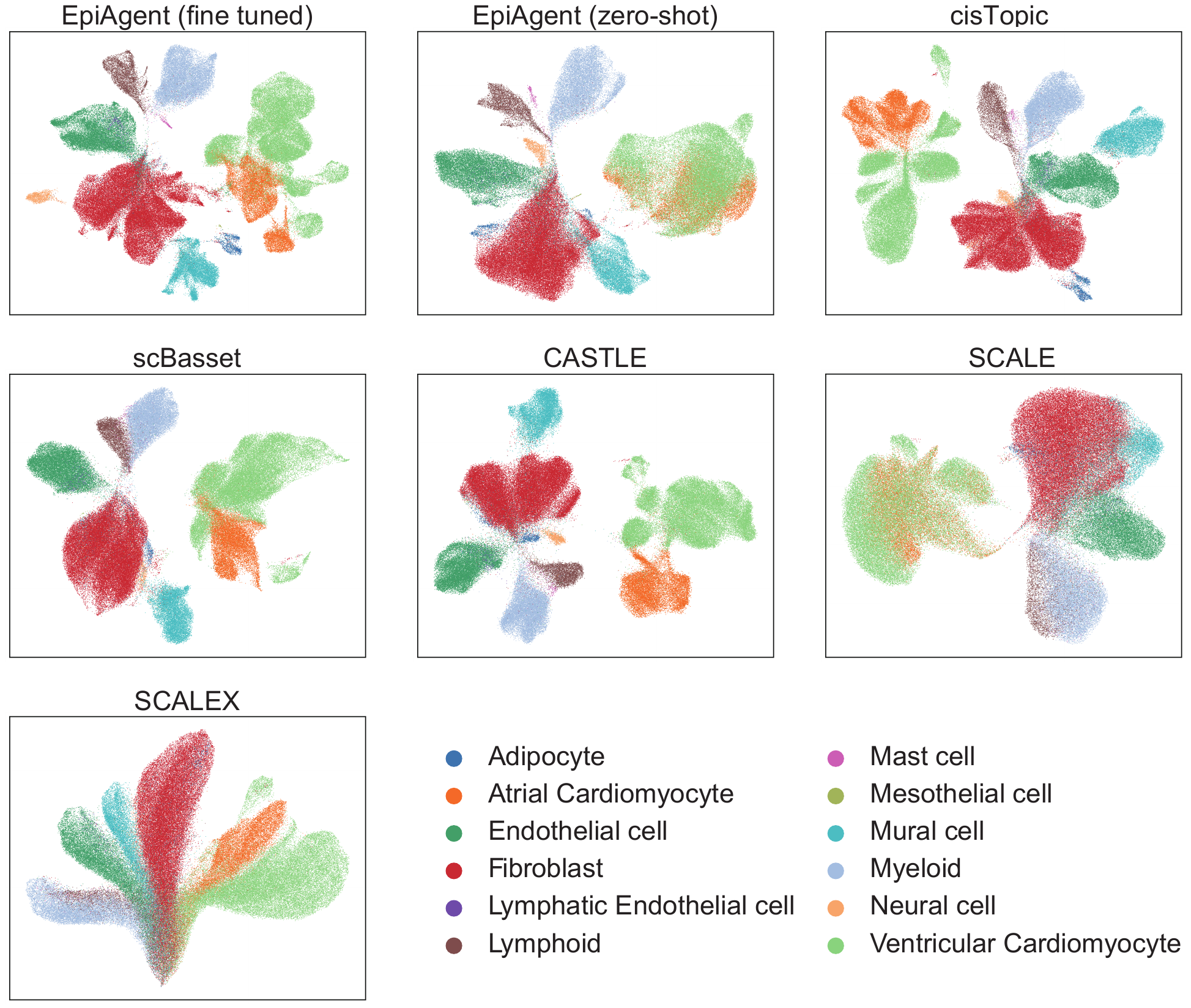

**Supplementary Figure 2.** UMAP visualization of cell embeddings derived from different methods on the Kanemaru2023 dataset.
**Supplementaery Figure 3**

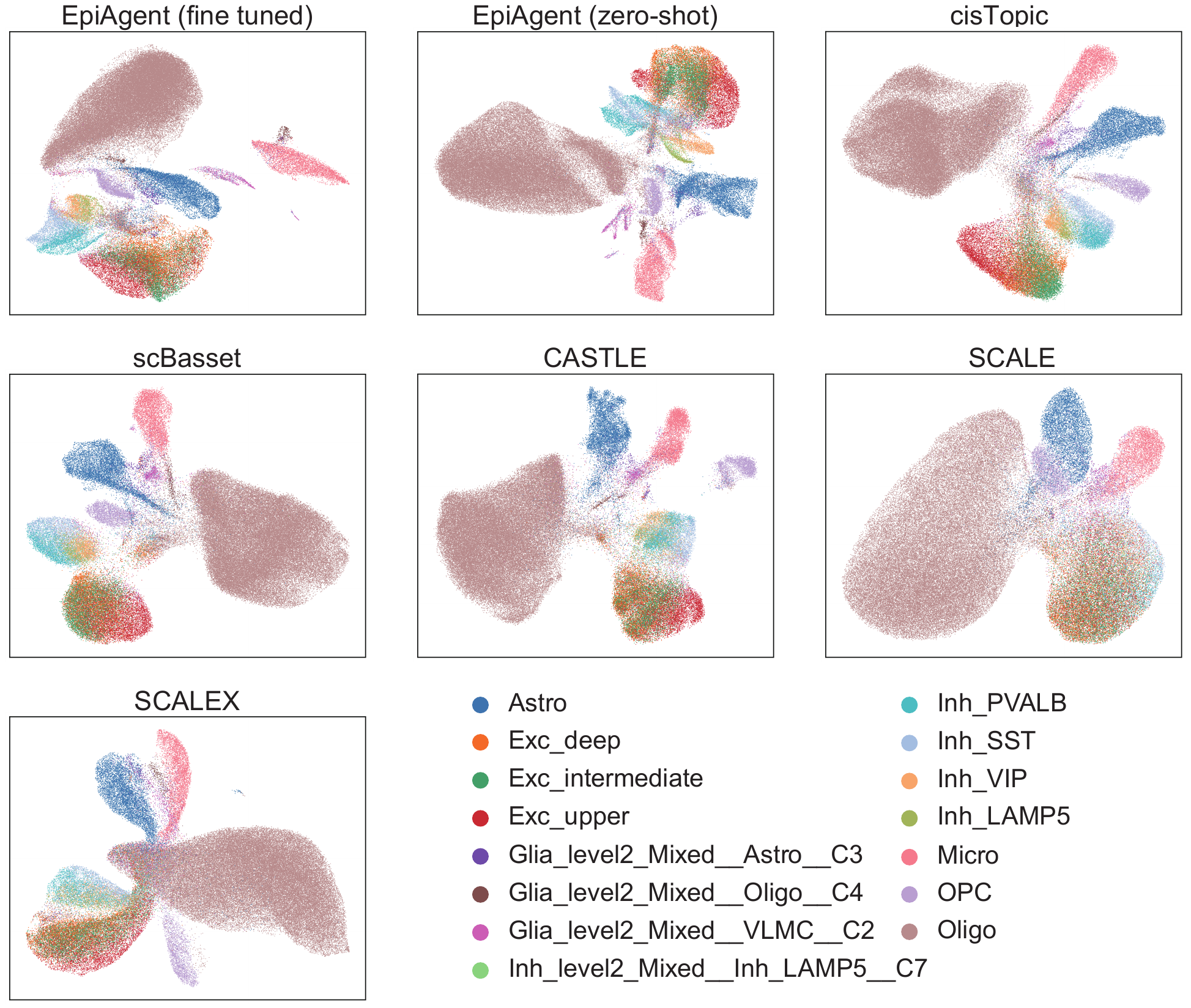

**Supplementary Figure 3.** UMAP visualization of cell embeddings derived from different methods on the Li2023b dataset.

**Supplementary Figure 4**

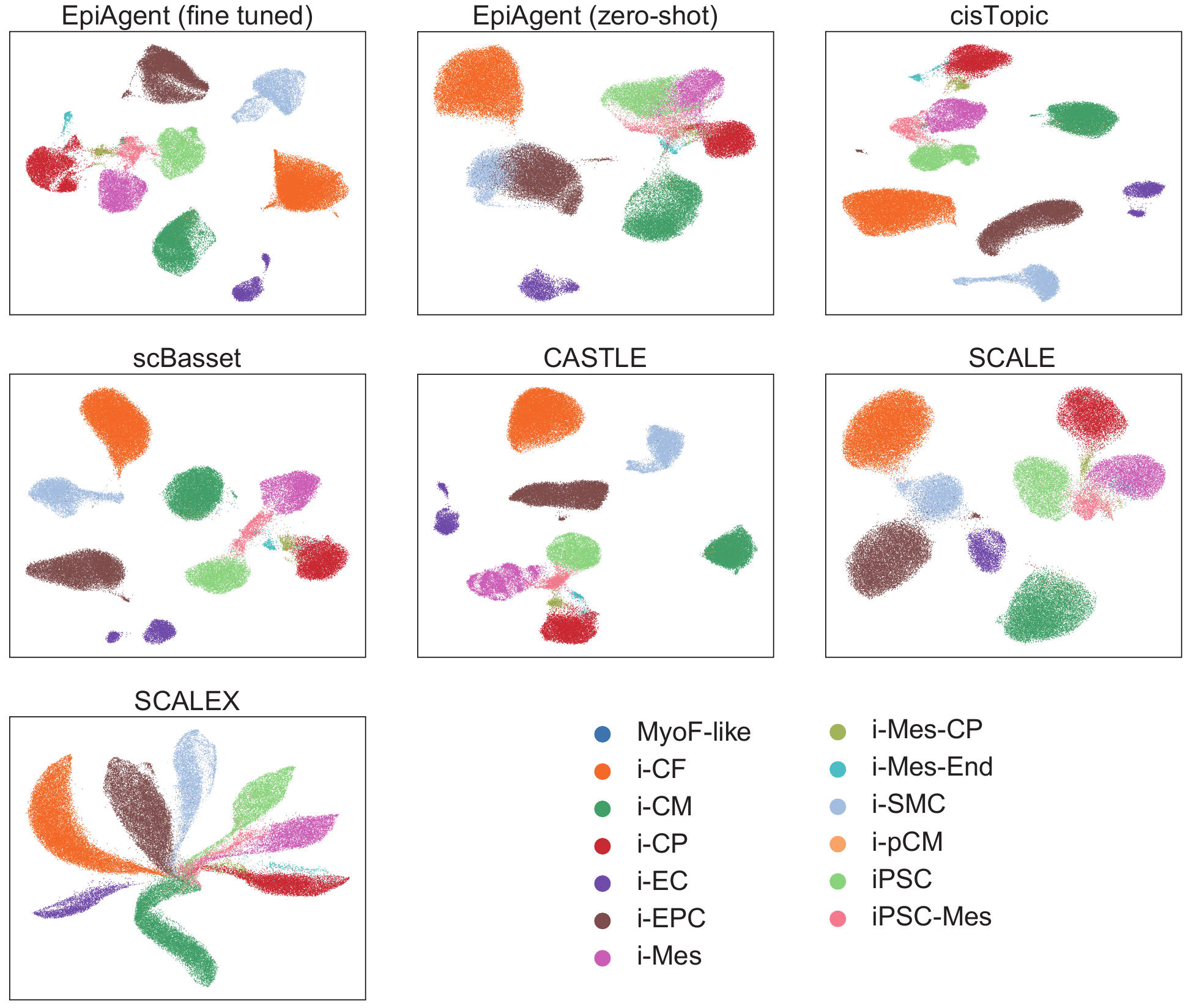

**Supplementary Figure 4.** UMAP visualization of cell embeddings derived from different methods on the Ameen2022 dataset.

**Supplementary Figure 5**

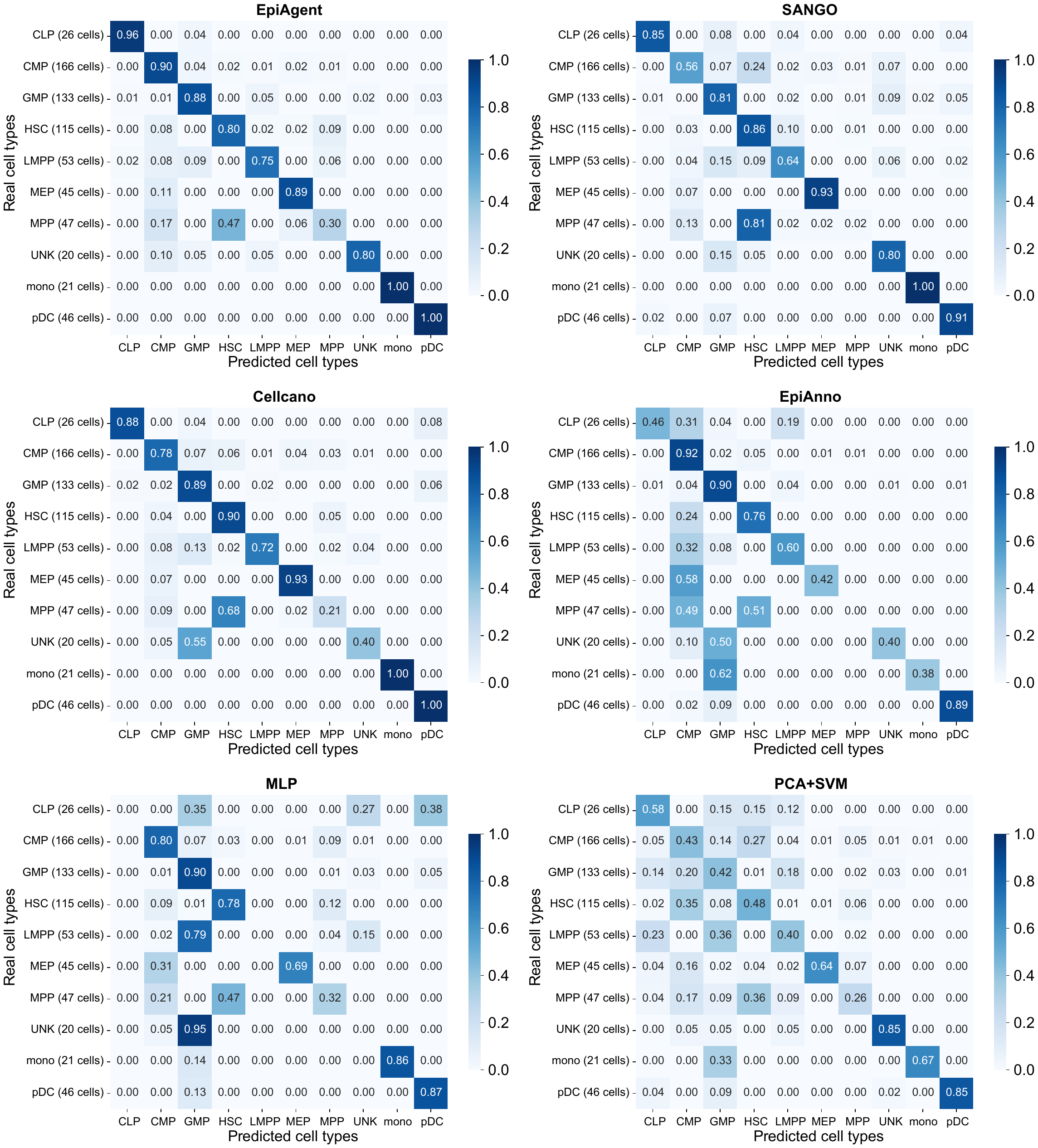

**Supplementary Figure 5.** Confusion matrix derived from different methods on the Buenrostro2018 dataset. Each dataset is randomly split, with two-thirds serving as the reference dataset for training and the remaining one-third as the query dataset. The numbers in parentheses next to each label on the Y-axis indicate the count of cells for that cell type in the query dataset.

**Supplementary Figure 6**

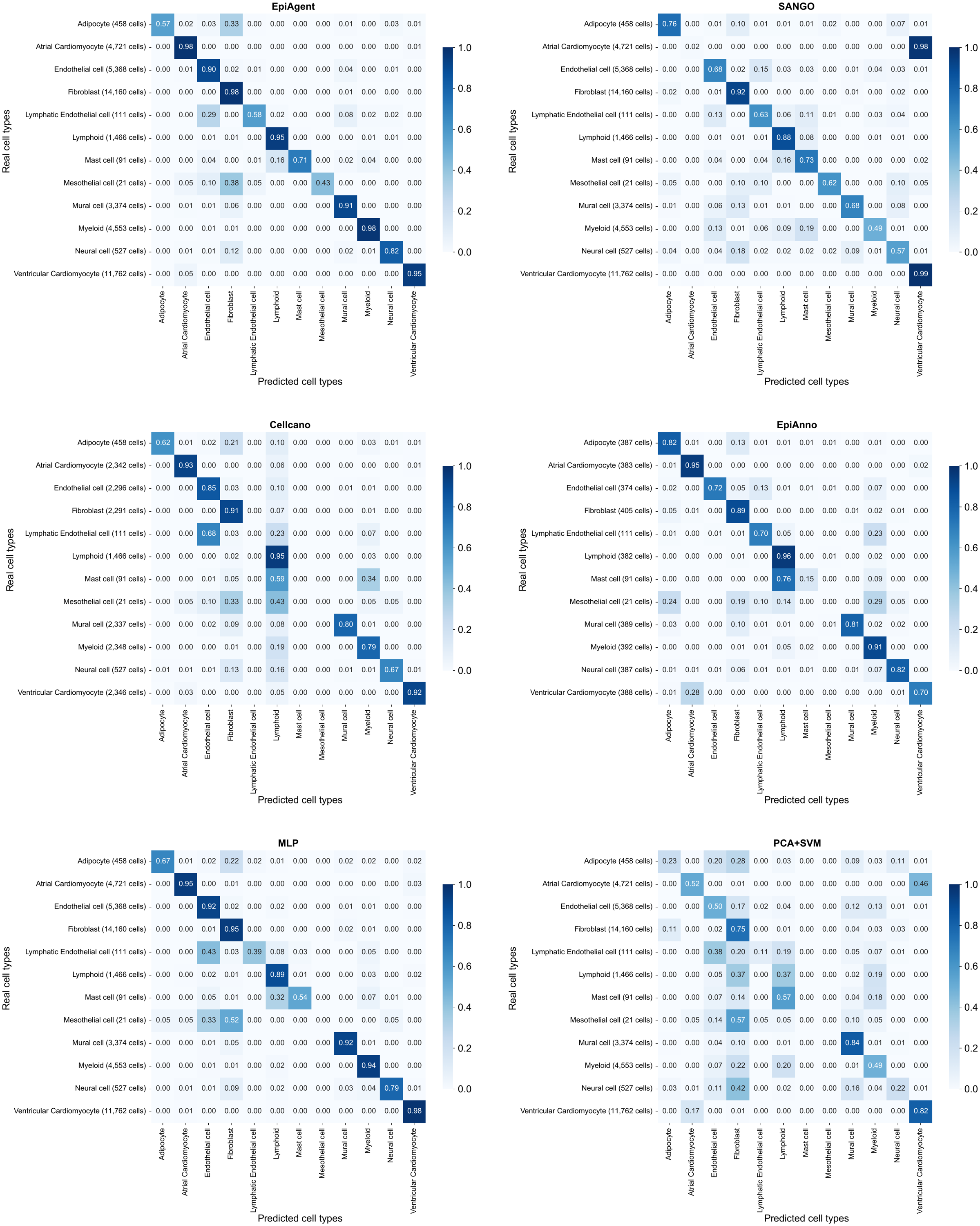

**Supplementary Figure 6.** Confusion matrix derived from different methods on the Kanemaru2023 dataset. Each dataset is randomly split, with two-thirds serving as the reference dataset for training and the remaining one-third as the query dataset. The numbers in parentheses next to each label on the Y-axis indicate the count of cells for that cell type in the query dataset.

**Supplementary Figure 7**

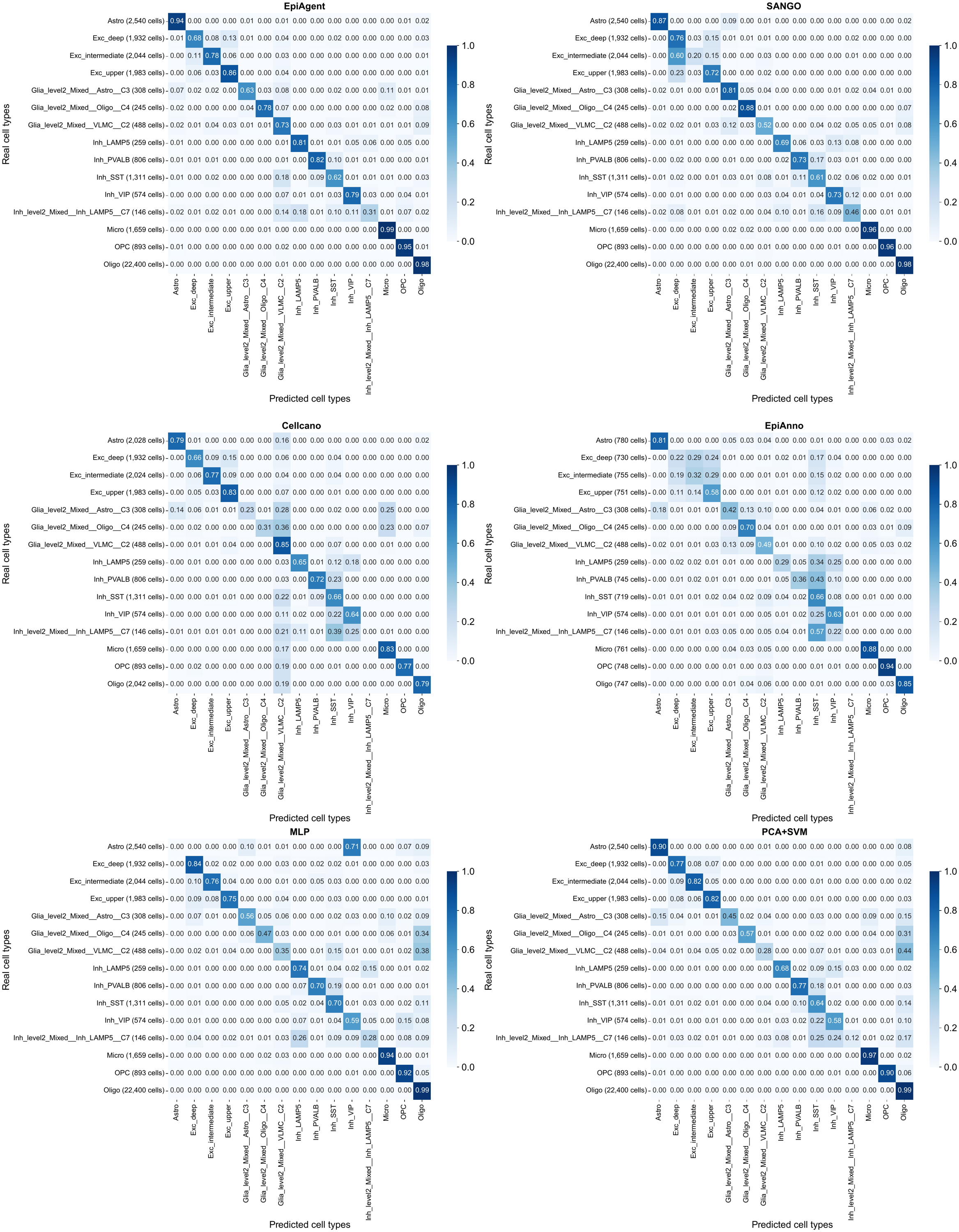

**Supplementary Figure 7.** Confusion matrix derived from different methods on the Li2023b dataset. Each dataset is randomly split, with two-thirds serving as the reference dataset for training and the remaining one-third as the query dataset. The numbers in parentheses next to each label on the Y-axis indicate the count of cells for that cell type in the query dataset.

**Supplementary Figure 8**

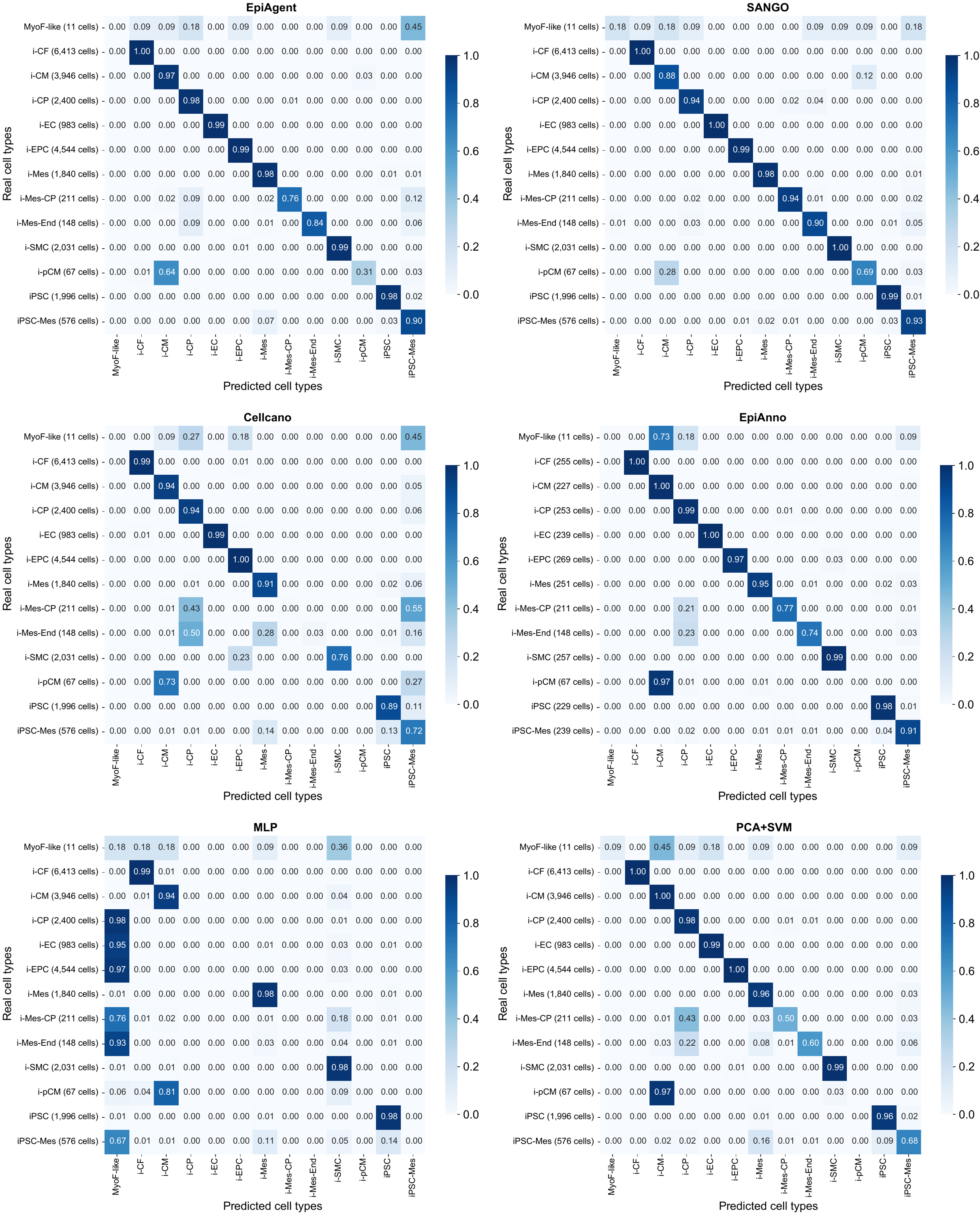

**Supplementary Figure 8.** Confusion matrix derived from different methods on the Ameen2022 dataset. Each dataset is randomly split, with two-thirds serving as the reference dataset for training and the remaining one-third as the query dataset. The numbers in parentheses next to each label on the Y-axis indicate the count of cells for that cell type in the query dataset.

**Supplementary Figure 9**

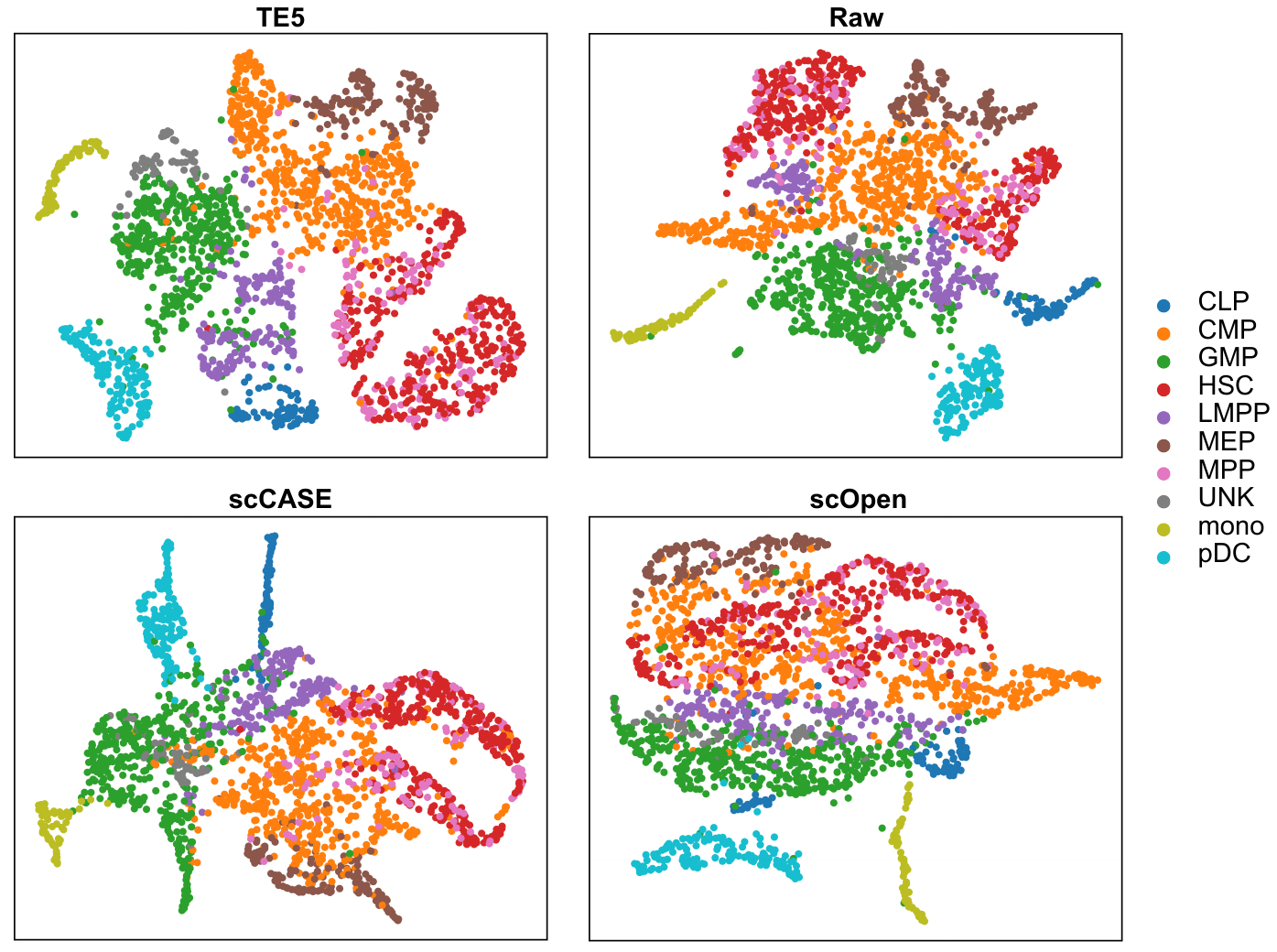

**Supplementary Figure 9.** UMAP visualization of PCA embeddings generated from imputed cell-by-cCRE matrices using various methods, compared to the PCA embeddings derived from the raw cell-by-cCRE matrix, on the Buenrostro2018 dataset.

**Supplementary Figure 10**

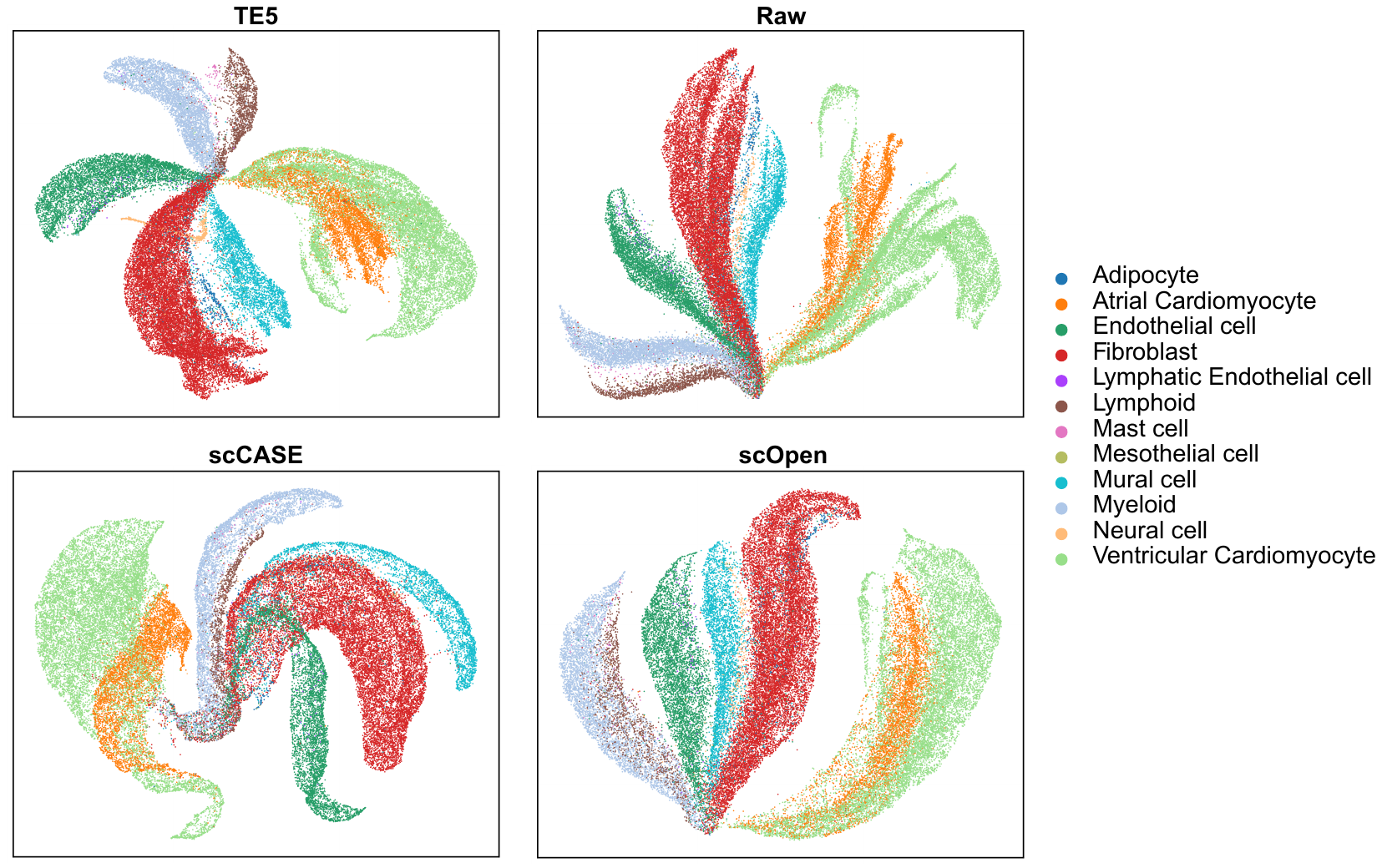

**Supplementary Figure 10.** UMAP visualization of PCA embeddings generated from imputed cell-by-cCRE matrices using various methods, compared to the PCA embeddings derived from the raw cell-by-cCRE matrix, on the Kanemaru2023 dataset.

**Supplementary Figure 11**

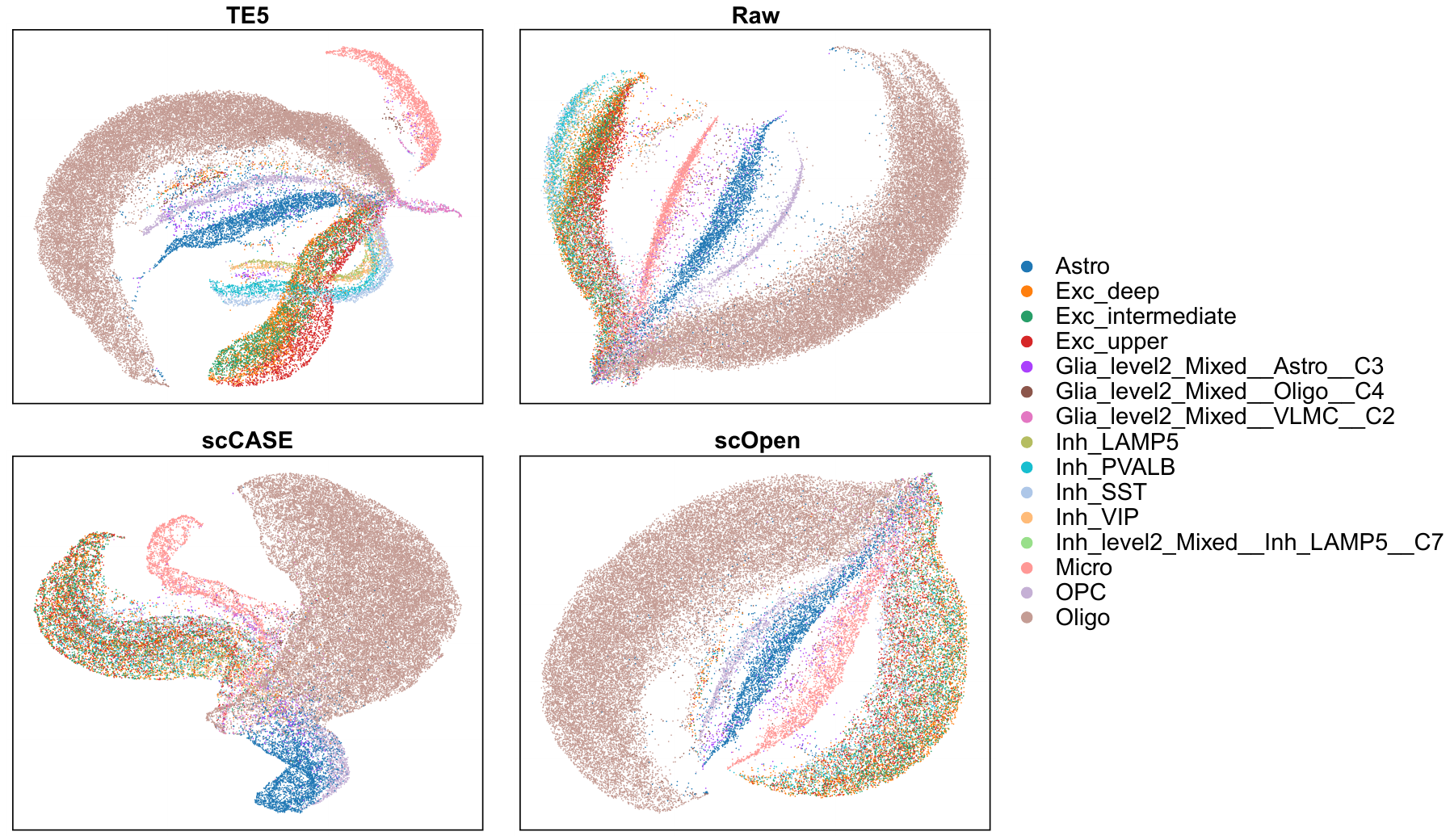

**Supplementary Figure 11.** UMAP visualization of PCA embeddings generated from imputed cell-by-cCRE matrices using various methods, compared to the PCA embeddings derived from the raw cell-by-cCRE matrix, on the Li2023b dataset.

**Supplementary Figure 12**

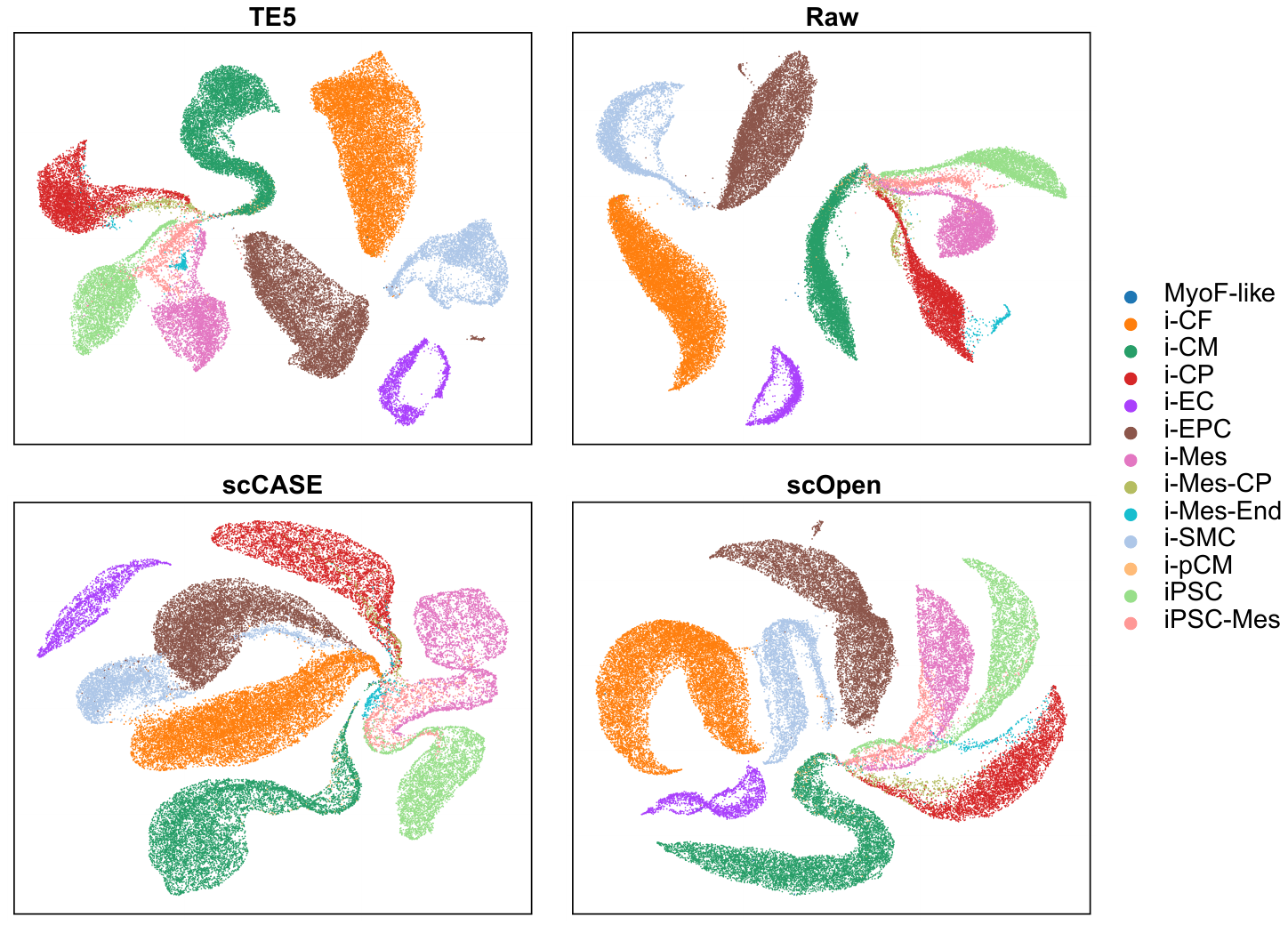

**Supplementary Figure 12.** UMAP visualization of PCA embeddings generated from imputed cell-by-cCRE matrices using various methods, compared to the PCA embeddings derived from the raw cell-by-cCRE matrix, on the Ameen2022 dataset.

**Supplementary Figure 13**

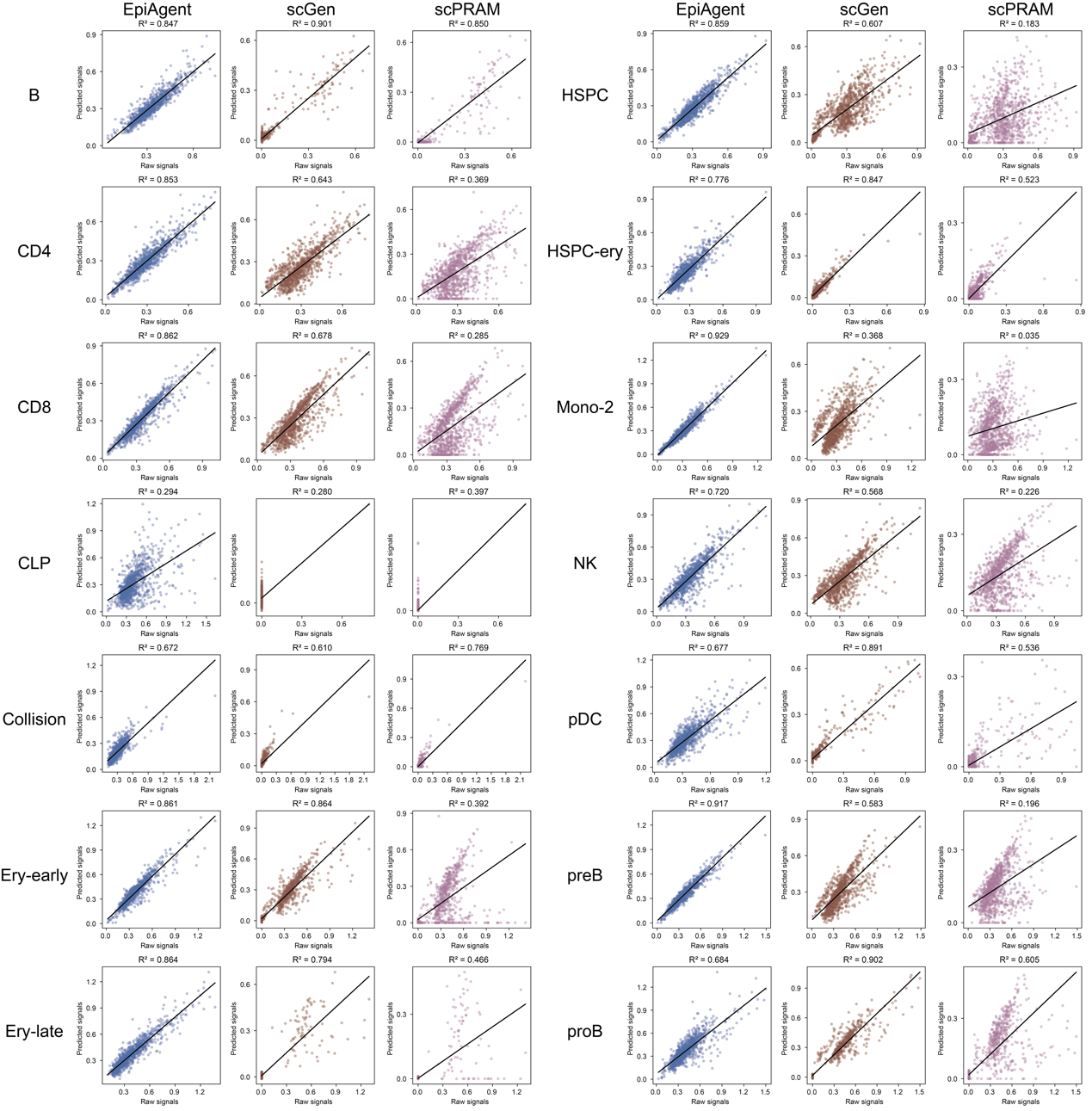

**Supplementary Figure 13.** Linear regression fit for mean accessibility of the top 1,000 DA cCREs between raw signals and predicted signals from different methods, with cells from various cell types serving as query data. Each column serves as results from a specific method, and each row serves as results with cells from a specific cell type serving as query data. The title of each subplot indicates the coefficient of determination (R²) for the linear regression fit.

**Supplementary Figure 14**

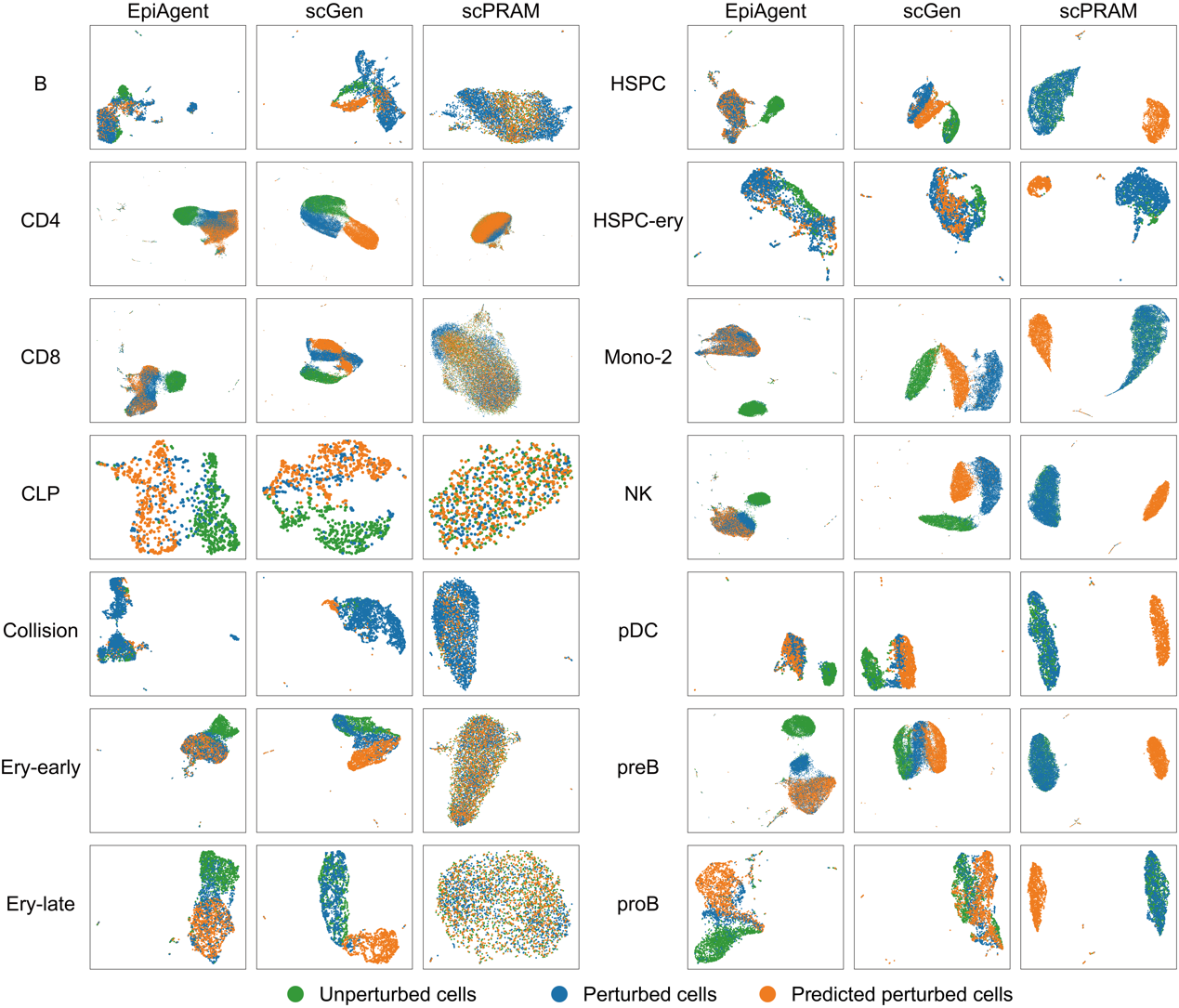

**Supplementary Figure 14.** UMAP visualization of embeddings for unperturbed cells, perturbed cells and predicted perturbed cells, derived from different methods. Each column serves as results from a specific method, and each row serves as results with cells from a specific cell type serving as query data.

**Supplementary Figure 15**

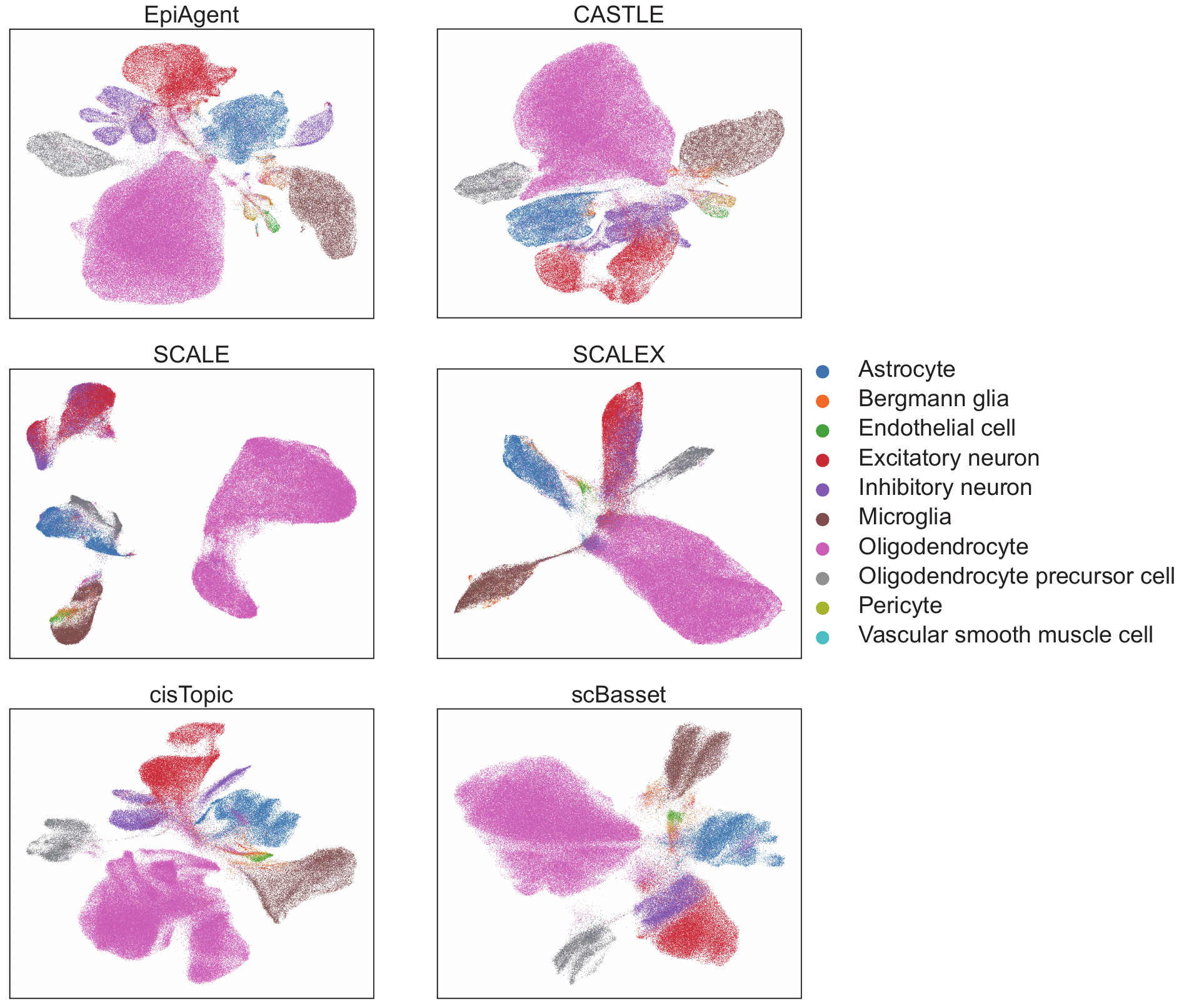

**Supplementary Figure 15.** UMAP visualization of cell embeddings derived from different methods on the Li2023a, Lee2023 and Li2023b datasets, using cell types as labels.

**Supplementary Figure 16**

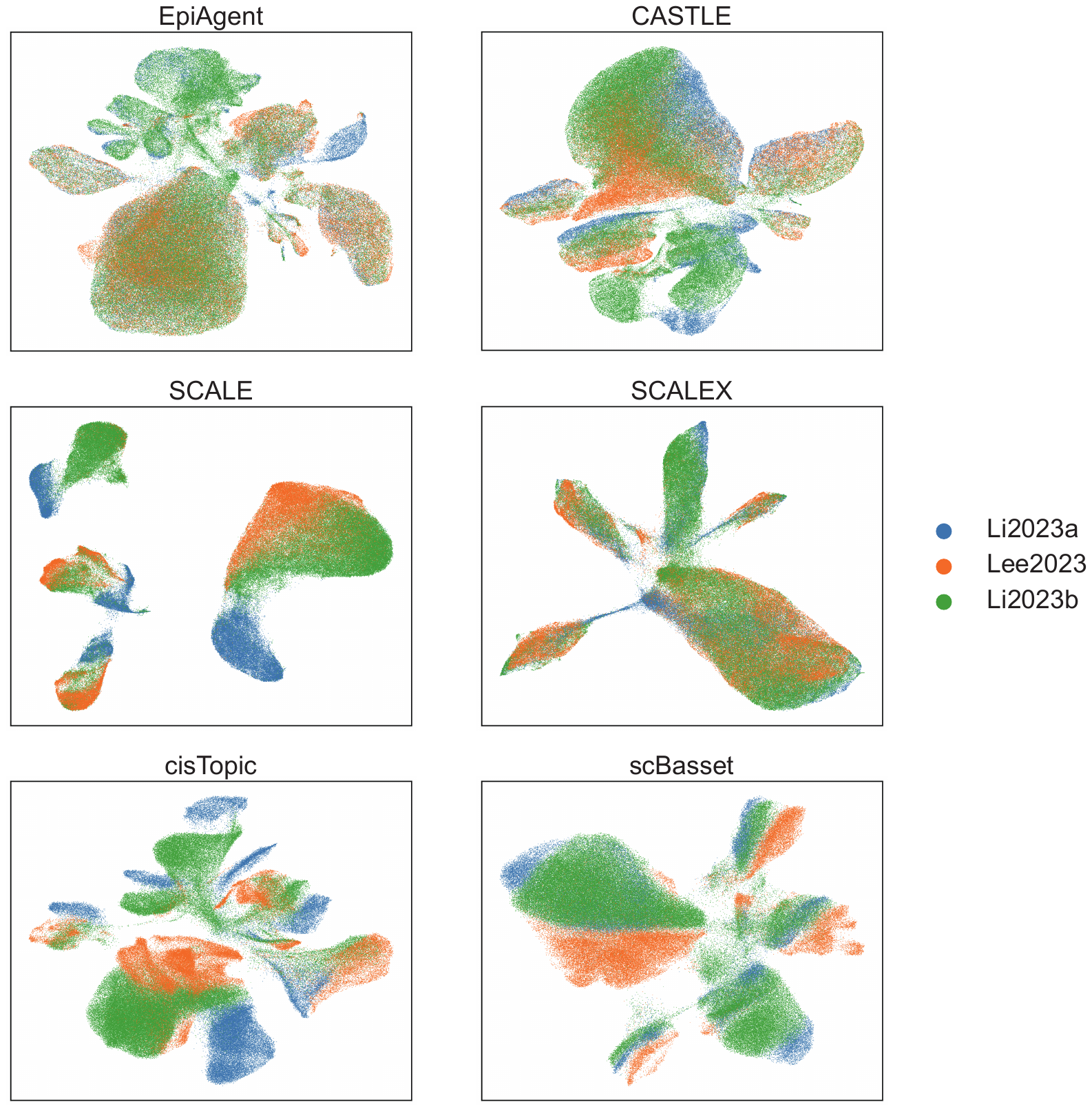

**Supplementary Figure 16.** UMAP visualization of cell embeddings derived from different methods on the Li2023a, Lee2023 and Li2023b datasets, using batches (datasets) as labels.

**Supplementary Figure 17**

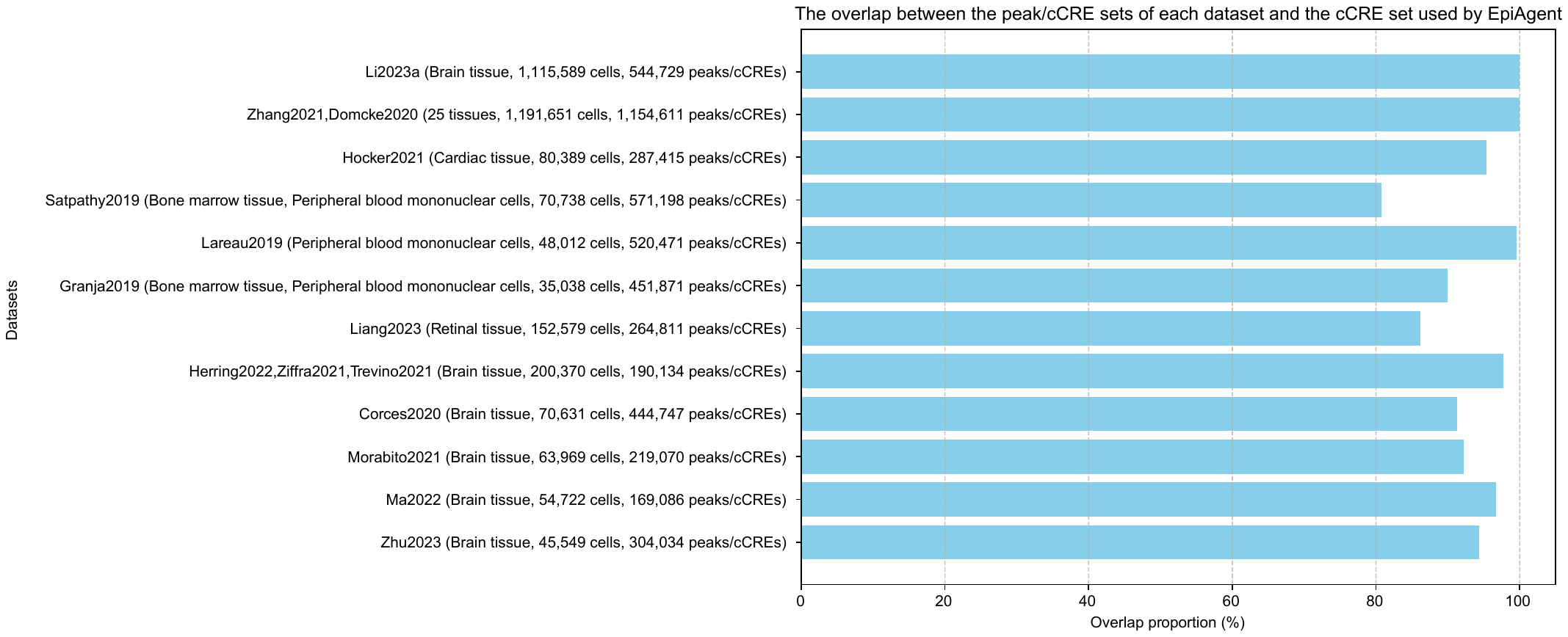

**Supplementary Figure 17.** The overlap proportion (%) between the peak/cCRE sets of each dataset and the cCRE set used by EpiAgent.

**Supplementary Figure 18**

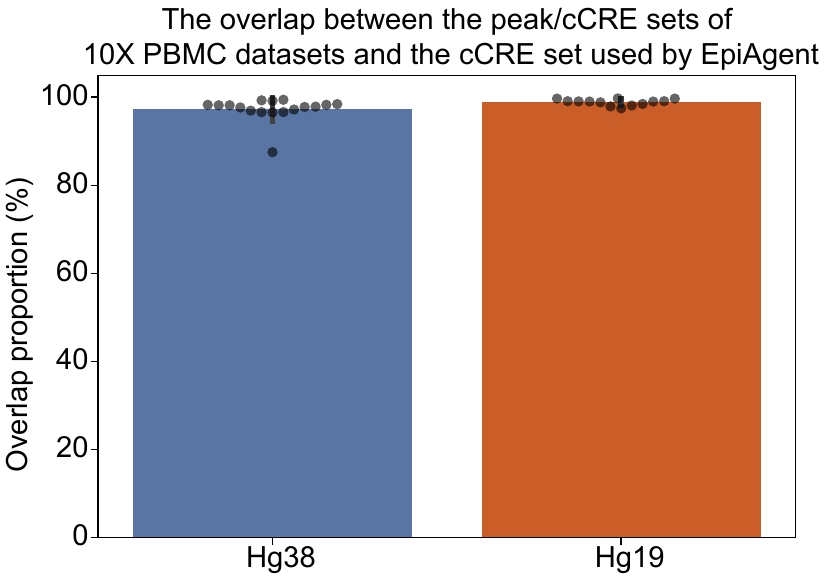

**Supplementary Figure 18.** The overlap proportion (%) between the peak/cCRE sets of 10X PBMC datasets splited by reference genome and the cCRE set used by EpiAgent.

**Supplementary Figure 19**

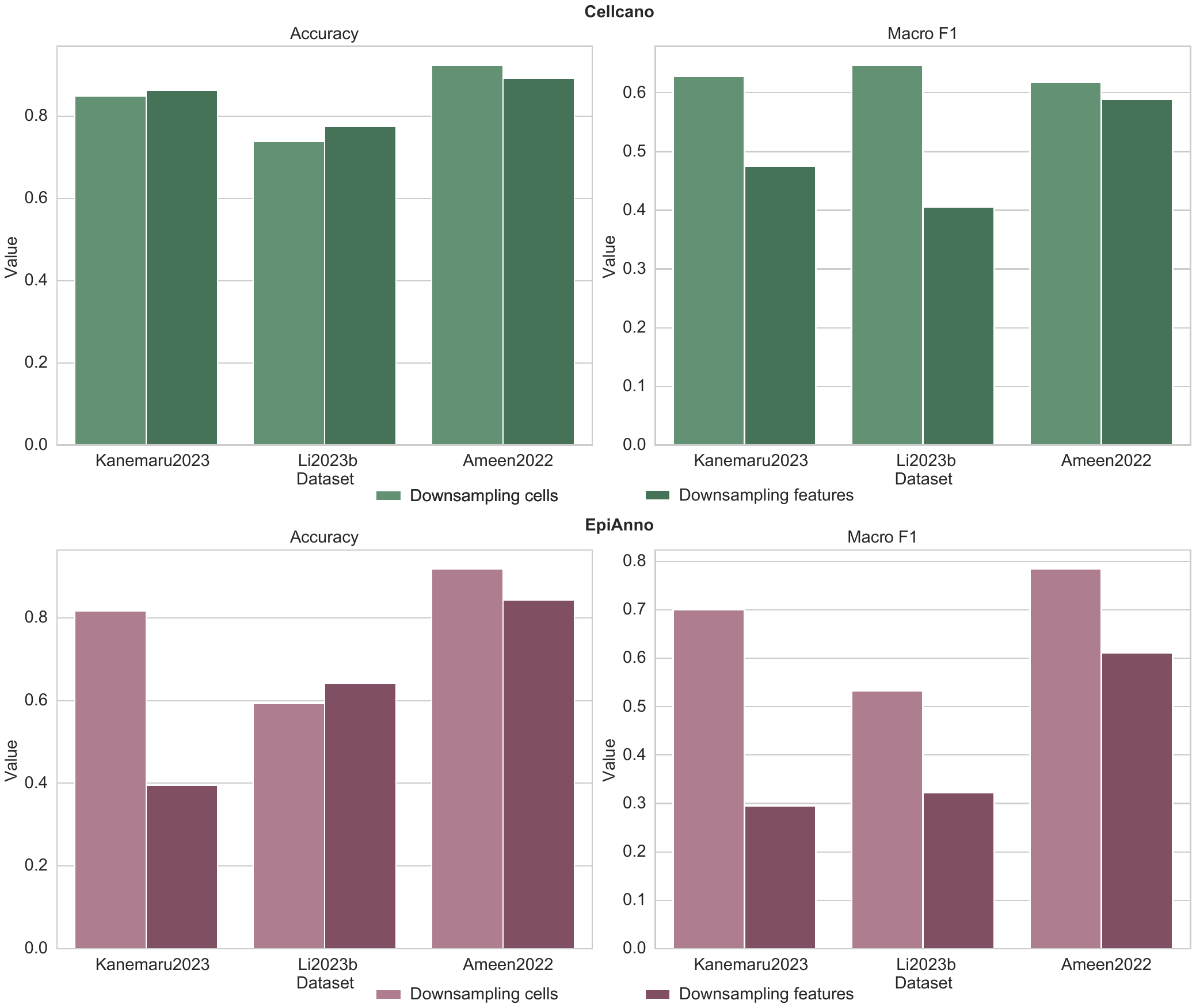

**Supplementary Figure 19.** Performance comparison of Cellcano and EpiAnno under two downsampling strategies: downsampling cells and downsampling features, evaluated using accuracy and macro F1 scores. The downsampling cells strategy involves randomly reducing the number of cells, while the downsampling features strategy involves randomly reducing the number of features. In both strategies, downsampling is performed iteratively until methods can run successfully under the reduced parameters.
